# RSV Infects the Human Nasal Epithelium via the Basolateral Route with Distinct Subgroup Infectivity and Basal Cell Tropism

**DOI:** 10.64898/2026.03.05.709813

**Authors:** Ashley Murray, Divya Nagaraj, Emily M. Schultz, Gina Aloisio, Erin Nicholson, Sarah E. Blutt, Vasanthi Avadhanula, Pedro A. Piedra

## Abstract

Respiratory syncytial virus (RSV) causes millions of lower respiratory tract infections (LRTIs) in young children, older adults, and immunocompromised populations every year. RSV infection initiates in the upper respiratory tract and can progress to the lower airways resulting in bronchiolitis, pneumonia, and even death. RSV primarily infects epithelial cells apically, but we hypothesized that basolateral exposure of the respiratory epithelium could provide an alternative mechanism of infection that contributes to LRTI development. Using a human nose organoid-air liquid interface (HNO-ALI) model, we performed apical and basolateral inoculations with contemporaneous RSV strains (RSV/A/ON and RSV/B/BA) representing the two RSV subgroups (A and B) in both adult and infant derived HNO-ALIs. Basolateral RSV exposure resulted in delayed viral replication and apical release compared to apical infection. A statistically significant difference in basolateral infection frequency was observed between RSV/B/BA and RSV/A/ON (81.3% versus 25%). Basolateral infection selectively targeted a rare basal cell population, while preserving epithelial integrity. Using undifferentiated HNO-ALIs, we determined for the first time that Krt23+ activated basal cells (ABCs) are uniquely susceptible to RSV infection, a finding we confirmed in fully differentiated HNO-ALIs. Together, our findings show that RSV can infect the respiratory epithelium from the basolateral side by initially targeting a rare subset of basal cells before spreading apically to ciliated cells. Moreover, RSV/B/BA may have an advantage over RSV/A/ON in utilizing the basolateral infection route. These findings highlight an alternative RSV infection pathway and could be a potential mechanism for RSV spread to the lower airways.

**Importance:** Understanding the pathogenesis of RSV is essential to understanding and preventing acute and long-term sequelae from infection. The canonical understanding of RSV infection is that the virus infects and is restricted to the apical ciliated cells upon inhalation or fomite exposure. We demonstrate that an alternative route of infection – the basolateral route, can be utilized by RSV to infect the apical ciliated cells of the respiratory epithelium. We also show for the first time a novel difference in infectivity between the two contemporaneous RSV strains (RSV/A/ON and RSV/B/BA). In addition, we describe a rare basal subset-the Krt23+ activated basal cells that are uniquely susceptible to RSV thus expanding the known cellular tropism of RSV. Infection of basal cells can impact airway differentiation, homeostasis, and remodeling. Overall, our findings expand on the pathogenesis of RSV and indicate there are alternative mechanisms of infection and cell populations that are susceptible to RSV.

## Introduction

RSV is one of the leading causes of lower respiratory tract infections (LRTIs), causing approximately 33.8 million cases, 3.4 million hospitalizations, and 66,000-199,000 deaths globally in children <5 years old (1). In addition, RSV LRTIs are a source of morbidity and mortality amongst older adults, immunocompromised populations, and those with comorbidities (2, 3). Despite the global burden caused by RSV LRTIs, it remains unclear as to how RSV can progress from a self-limited upper respiratory tract infection (URTI) to a severe and life-threatening LRTI. The main hypothesis for RSV dissemination to the lower airways suggests the aspiration of sloughed off RSV infected epithelial cells and/or aspirated viral particles to the lower airways to be responsible for LRTI development (4). However, the specific mechanism of spread by RSV from upper to lower airways has yet to be characterized. An alternative hypothesis suggests that during an RSV URTI, epithelial damage and loss of barrier integrity due to the inflammatory immune response and cell death could allow for viral spread from the upper respiratory tract and contribute to the development of RSV LRTI (5, 6). Several reports have described RSV dissemination to extrapulmonary sites and detection of viral RNA in peripheral blood mononuclear cells (PBMCs) (7–10). Additionally, *Torres et al*. reported a correlation between RSV RNA in peripheral blood and disease severity in mice (11). Taken together, these reports suggest RSV’s ability to initiate infection of the lower respiratory tract via an alternative route of infection, but to date this remains uninvestigated.

We previously reported the use of the clinically and physiologically relevant human nose organoid – air liquid interface (HNO-ALI) model to study the pathogenesis of RSV (12, 13). HNO-ALIs are comprised of all the major cell types found within the polarized pseudostratified respiratory epithelium such as ciliated, goblet, club, and basal cells (13). Ciliated cells, the major cell type on the apical surface, are responsible for mucociliary clearance of pathogens and cellular debris while secretory cells like goblet and club cells are the main secretors of mucus and protective substances, respectively (14). Basal cells, found at the base of the epithelium, are the progenitor stem cells that differentiate into other epithelial cells and are responsible for epithelial regeneration (14). The HNO-ALI model is susceptible to RSV infection and generates an innate immune epithelial response to both RSV and SARS-CoV-2 (12, 13, 15). The primary target cell population of RSV is the apical ciliated cells of the respiratory epithelium(16). However, Persson *et al*. previously reported that basal cells are susceptible to RSV infection in a scratch injury model using differentiated human bronchial epithelial cells (17). Using our HNO-ALI model, we sought to understand if RSV can utilize the basolateral route to infect the respiratory epithelium.

In this study, we characterized the basolateral route of RSV infection by inoculating the basolateral compartment of HNO-ALIs with contemporaneous strains RSV/A/Ontario (RSV/A/ON) or RSV/B/Buenos Aires (RSV/B/BA) of the two RSV subgroups – RSV/A and RSV/B. We found that the basolateral route of exposure is a viable infection route that leads to infection of the apical ciliated cells with apical release of the virus. However, there was a surprising difference in infection outcome between the two RSV strains with RSV/B/BA having a statistically significant higher rate of infection compared to RSV/A/ON. Additionally, we identified a specific subset of rare basal cells (Krt23+ activated basal cells) that are susceptible to RSV infection. Taken together, our findings show for the first time that under basolateral exposure conditions, RSV can infect the respiratory epithelium by first infecting rare Krt23+ activated basal cells, with subsequent spread to the apical ciliated cells, potentially demonstrating an alternate route of spread to the lower respiratory tract.

## Results

### RSV infects the respiratory epithelium from the basolateral side and infection outcome varies between contemporaneous RSV strains of both subgroups

To determine if the basolateral route of infection was a viable route of infection, eight adult-derived and eight infant-derived HNO-ALI lines were basolaterally inoculated with contemporaneous RSV/A/ON and RSV/B/BA. The age and sex of the various donors are represented in Supplemental Table 1. Preliminary infection studies determined that a high MOI of 1.0 was needed for the basolateral route of infection (Supplemental Figure 1A-B), in contrast to apical infection where a low MOI of 0.01 is sufficient for productive infection. (12, 13).

In both adult and infant HNO-ALIs, RSV infected through the basolateral route of infection, resulting in the apical release of infectious virus between 5 to 8 days post inoculation (dpi) as represented in three adult and three infant HNO-ALI lines (Figure 1A-D). qPCR of basolateral media and apical wash samples showed steady levels of viral RNA detected in the basolateral compartment over time and delayed but incremental increase in the amount of viral RNA detected on the apical side, demonstrating an eclipse phase of 2 to 8 days from initial inoculation on the basolateral side to apical release of the virus (Supplemental Figure 2A-D). Interestingly, there was variability in time until apical release of the virus between transwell replicates from the same donor line as well as across different donor lines.

**Figure 1:**
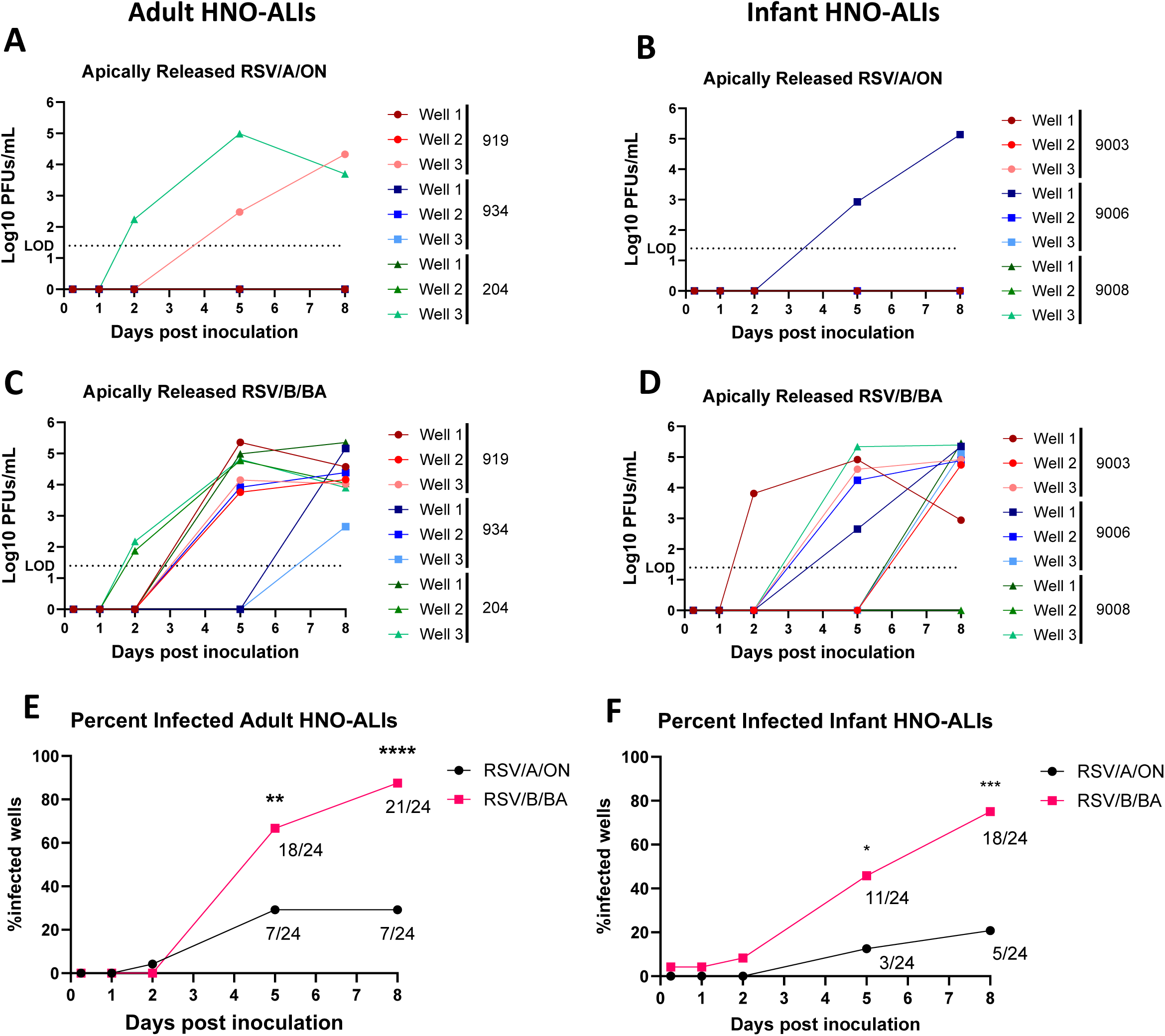
Infectious RSV is apically released with basolateral inoculation, but infection outcome differs between subgroups. Apically released plaque forming units of RSV/A/ON in (A) adult and (B) in infant HNO-ALI cultures. Apically released plaque forming units of RSV/B/BA in (C) adult and (D) infant HNO-ALIs. Data derived from triplicate transwell replicates from three representative adult donor lines (919, 934, 204) and infant donor lines (9003, 9006, 9008) are shown. LOD = limit of detection. Total percentage of HNO-ALI transwells that demonstrated apical release of RSV with basolateral inoculation in (E) adult (n=8 lines, triplicate wells/line) and in (F) infant (n=8 lines, triplicate wells/line). Statistical significance determined by Fisher’s Exact test; p values: *<0,05; **<0.01; *** <0.001; **** <0.0001.

Unexpectedly, we observed that RSV/B/BA infected at a greater frequency compared to RSV/A/ON (Figure 1A-D). Plaque assay data from a subset of three representative adult HNO-ALI lines with triplicate technical replicates (total 9 transwells) demonstrated RSV/A/ON infected 2/9 transwells (Figure 1A) while RSV/B/BA infected 9/9 (Figure 1C). Similarly, plaque assay data from a subset of three representative infant HNO-ALI lines with triplicate technical replicates (total 9 transwells) demonstrated RSV/A/ON infected 1/9 transwells (Figure 1B) and RSV/B/BA infected 8/9 (Figure 1D).

Using apically released infectious virus as a proxy for infection status, we next evaluated the complete data set of all eight adults and eight infant HNO-ALIs lines. RSV/B/BA infected a statistically significantly greater proportion of HNO-ALIs across eight adult (Figure 1E) and eight infant (Figure 1F) donor lines. 87.5% of adult-derived and 75% of infant-derived HNO-ALIs were infected by RSV/B/BA at 8 dpi. In contrast, RSV/A/ON infected only 29.2% of adult-derived and 20.8% of infant-derived HNO-ALIs at the same time point (Figure 1E–F). Collectively, RSV/B/BA infected 81.3% (39 of 48) of all HNO-ALI transwells, while RSV/A/ON infected 25% (12 of 48). Notably, we observed a difference in infection outcome between RSV/A/ON and RSV/B/BA in both technical replicates from the same donor and between donor lines. This variability between technical replicates and donor lines was specific to basolateral infection and has not been observed with apical infection and requires further study (12, 13). Together, these findings reveal a previously unrecognized difference in infectivity between two contemporaneous strains of different RSV subgroups, with RSV/B/BA demonstrating an enhanced capacity to initiate infection via the basolateral route in HNO-ALI cultures.

RSV primarily infects the respiratory epithelium via the apical route and in both our previous studies and the current work we observed no difference in infectivity in RSV subgroups using this traditional apical route of infection (12, 13). RSV/A/ON and RSV/B/BA strains did not show any difference in infectivity in four adult and four infant HNO-ALI lines (Supplemental Figure 3A-B) (12, 13). Collectively, these data highlight that the enhanced infectivity of RSV/B/BA is specific to basolateral exposure conditions, suggesting that RSV/B/BA is better adapted than RSV/A/ON to establish infection via the basolateral route.

To validate this difference in infectivity observed with the basolateral route of inoculation between contemporaneous strains of both RSV subgroups, we repeated the infection in four adult HNO-ALI lines with an additional contemporaneous RSV/A/ON and RSV/B/BA strains (RSV/A/ON-Hou and RSV/B/BA-IP respectively), isolated in 2022. Importantly, the statistically significant difference in infection outcome was retained with these additional virus strains, with RSV/B/BA-IP infecting 66.7% of transwells and RSV/A/ON-Hou infecting 16.7% of transwells at 8 dpi (Supplemental Figure 4A). When the data from all four contemporaneous strains (two RSV/A/ON and two RSV/B/BA strains) were combined across the same four adult HNO-ALI lines, (total 24 transwells), the disparity in basolateral infectivity between strains of the two subgroups became even more pronounced. (Supplemental Figure 4B).

To characterize the progression of infection following basolateral inoculation, HNO-ALI transwells from a representative adult donor line were basolaterally inoculated with RSV/B/BA. Viral RNA levels were subsequently measured in the basolateral media, epithelial layer, and apical lumen over the course of infection. qPCR of the basolateral media, lysed epithelial layer, and apical lumen from RSV/B/BA infected transwells showed at early timepoints high viral RNA in the basolateral compartment, low intracellular viral RNA (epithelial layer), and undetectable viral RNA in the apical lumen (Supplemental Figure 5). As the infection progressed, the levels of intracellular viral RNA increased until there was apical detection of viral RNA at 4 to 6 dpi. By 8 dpi, there were elevated levels of both intracellular and apically detected viral RNA. This demonstrated that RSV/B/BA initiates infection on the basolateral surface with delayed spread through the epithelium before apical release, possibly explaining the eclipse phase of 2-5 days with no RSV detection between initial inoculation and apical release.

### RSV is detectable in apical ciliated cells at 8 dpi after basolateral inoculation and causes epithelial damage

To determine how the basolateral route of RSV infection impacts the respiratory epithelium, adult and infant HNO-ALIs at 8 dpi were sectioned and stained to evaluate epithelial damage and RSV localization. Immunofluorescence (IF) microscopy revealed that HNO-ALIs that were basolaterally inoculated with RSV/A/ON or RSV/B/BA with no apical release of RSV, looked identical to mock infected HNO-ALIs (Figure 2A, Supplemental Figure 6A-B). Both showed an intact apical ciliated cell layer, a distinct basal cell (Krt5+) layer, and lack of viral proteins. This non-infected phenotype was observed more frequently with RSV/A/ON inoculation, given that infection occurred less often in this group compared to RSV/B/BA inoculation. However, when infection and apical release of the virus occurred with basolateral RSV/A/ON or RSV/B/BA inoculation, HNO-ALIs displayed epithelial damage, ciliary loss, and RSV infected apical ciliated cells (Figure 2B-C). As expected from the viral kinetics data (Figure 1), apical ciliated cells were infected with RSV at 8 dpi. However, we also observed rare instances of RSV/A/ON and RSV/B/BA infected cells within the basal cell layer (white arrows). Notably, these cells were not always Krt5+, despite residing in the basal cell layer (Figure 2B-C).

**Figure 2:**
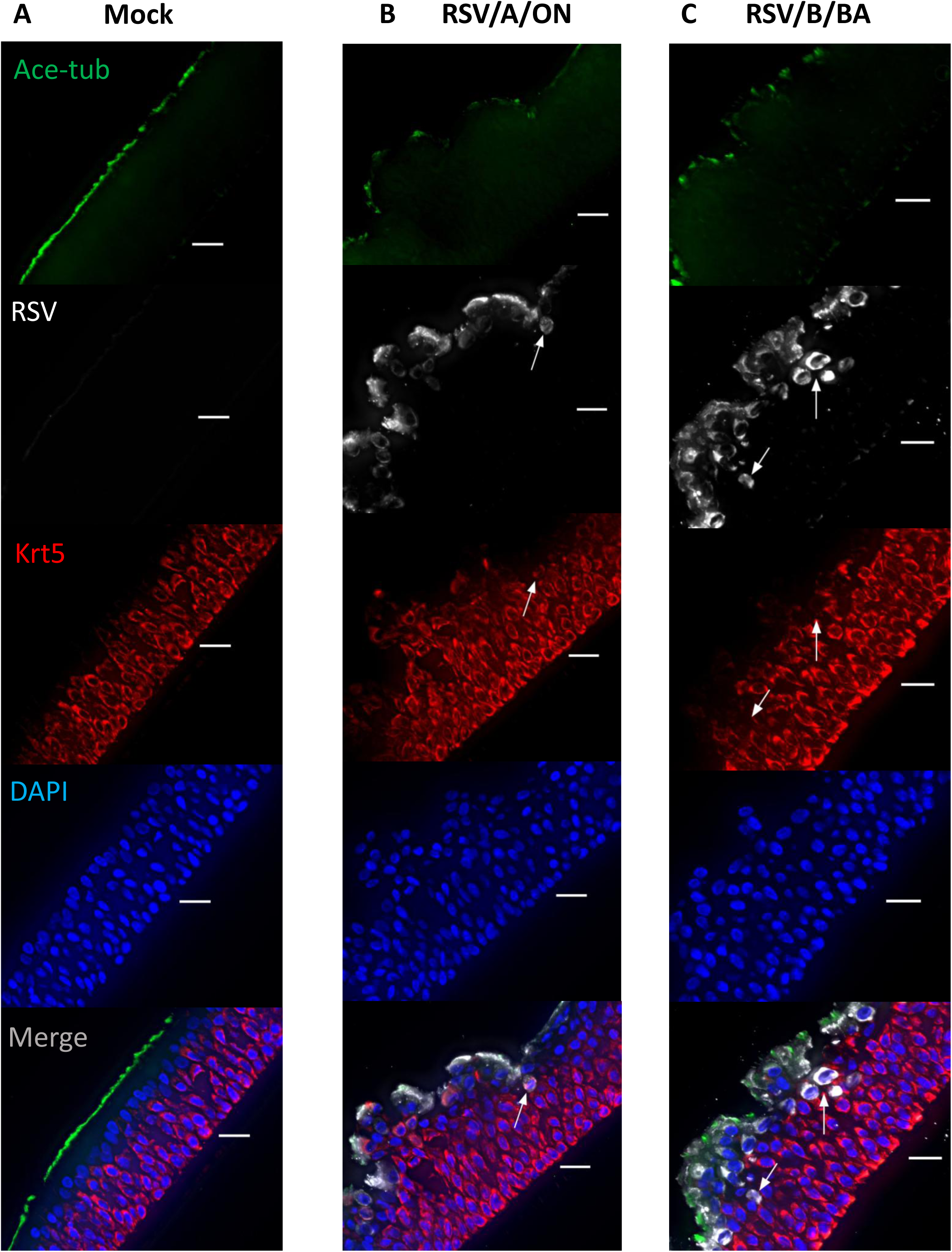
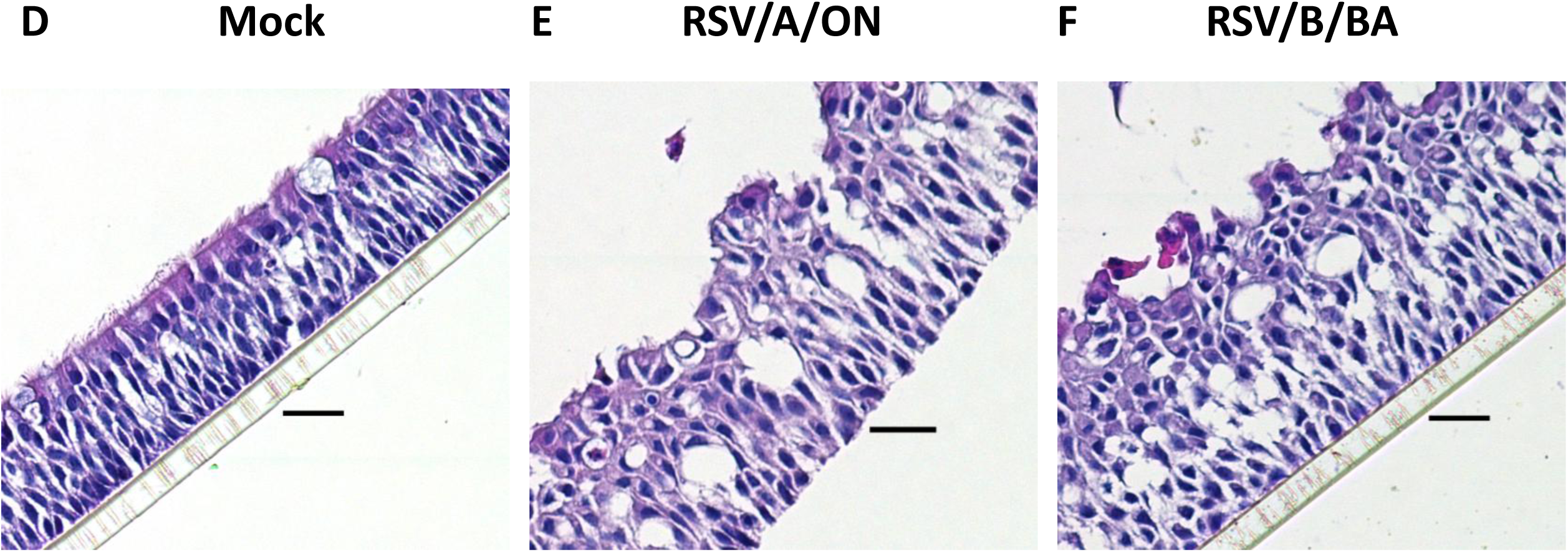
RSV detection and epithelial damage in HNO-ALIs with basolateral inoculation. Representative IF images of (A) mock, (B) RSV/A/ON, and (C) RSV/B/BA infected differentiated HNO-ALIs at 8 dpi from a single representative infant HNO-ALI line. Ciliated cells – Acetylated alpha-tubulin ‘Ace-tub’ (green), RSV (white), Basal cells – Krt5(red), and nuclei – DAPI (blue). Arrows denote RSV infected basal cells. Representative H&E images of (D) mock, (E) RSV/A/ON, and (F) RSV/B/BA infected HNO-ALIs at 8 dpi from a single representative infant HNO-ALI line. Scale bar is 20 µm.

H&E staining showed similar findings, in that a healthy, intact epithelium was observed in basolateral mock, RSV/A/ON, and RSV/B/BA inoculated HNO-ALIs with a non-infection phenotype (Figure 2D, Supplemental Figure 6C-D). Epithelial damage was observed in basolaterally inoculated RSV/A/ON and RSV/B/BA HNO-ALIs (Figure 2E-F) where apical infectious virus was detected (Figure 2B-C). Overall, successful basolateral RSV infection resulted in epithelial damage, delayed infection of ciliated cells, and rare instances of infected basal cells. Based on our initial findings we hypothesized that following basolateral inoculation, RSV will first infect an uncommon basal cell type before spreading to the apical ciliated cells where viral expansion and release would occur.

### TEER increases with RSV infection

We wanted to determine that apical release of RSV following basolateral inoculation was not due to loss of epithelial integrity, allowing the virus easy access to the apical ciliated cells. To rule out this possibility, adult and infant HNO-ALIs were apically or basolaterally inoculated with RSV/A/ON or RSV/B/BA. Simultaneous transepithelial electrical resistance (TEER) – a measure of epithelial integrity, and viral kinetics were monitored at each time point (Figure 3). qPCR of the apical wash samples from successful basolaterally infected wells showed apical detection of both RSV/A/ON and RSV/B/BA at 8 dpi (Data not shown). For the basolateral infection group, the average TEER for adult HNO-ALIs prior to infection was 1100 ohms (Figure 3A), while for infant HNO-ALIs the average TEER was 685 ohms (Figure 3B). Over the course of infection, TEER remained constant in mock-infected adult and infant HNO-ALIs. However, for successful RSV/A/ON and RSV/B/BA basolaterally infected HNO-ALIs, TEER increased during infection, with statistically significant differences in TEER seen at multiple timepoints compared to mock (Figure 3A-B). At 8 dpi, both RSV/A/ON and RSV/B/BA infected HNO-ALIs had significant differences in TEER compared to mock in adult and infant HNO-ALIs (Figure 3A-B). This result was mirrored with apical RSV infection, where again TEER remained constant in mock-infected HNO-ALIs, while RSV infection with strains from both RSV subgroups resulted in a statistically significant increase in TEER over time (Figure 3C-D). Overall, there was no early decrease in TEER that would indicate a loss of epithelial integrity with basolateral RSV inoculation that could contribute to translocation of the virus. Additionally, increased TEER was observed with basolateral and apical RSV infection, but to a greater degree with apical infection.

**Figure 3:**
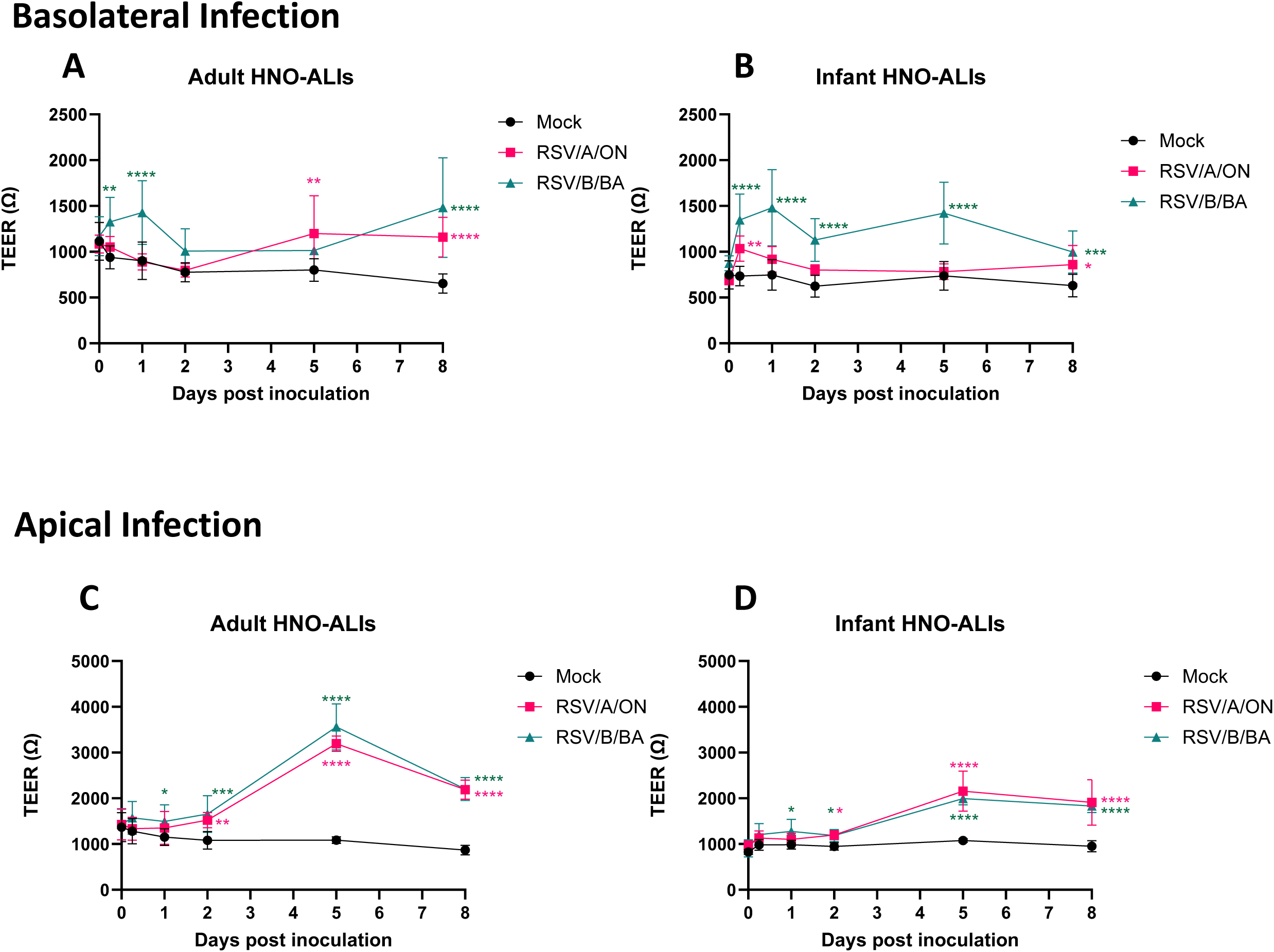
Changes in transepithelial electrical resistance (TEER) with both basolateral and apical RSV inoculation. TEER values in ohms over time with (A-B) basolateral and (C-D) apical RSV/A/ON and RSV/B/BA inoculation in differentiated adult and infant HNO-ALIs. Data represent mean ± standard deviation pooled from two adult and two infant lines with duplicate transwells/line and duplicate TEER measurements/transwell. Statistical significance determined by two-way ANOVA with Dunnett’s correction for multiple comparisons; p values: *<0.05; ***<0.001****<0.0001. Asterisk color indicates which virus had statistical significance at a particular timepoint.

Despite seeing TEER values increase with both apical and basolateral RSV infection, there was still clear damage to the apical side of the epithelium as observed in Figure 2, indicating that changes in epithelial integrity might not indicate a lack of damage due to viral infection, but more so a lack of epithelial breach.

### RSV infects a small subset of basal cells prior to delayed spread of the virus to the apical ciliated cell population

Since few RSV-infected basal cells were observed in basolateral inoculated HNO-ALIs at 8 dpi, we sought to investigate the delayed spread of RSV from basal to ciliated cells during basolateral inoculation. To do this, we used a sucrose purified recombinant GFP RSV/A2 (GFP-rRSV/A2), a prototypic genotype, to apically or basolaterally inoculate HNO-ALIs and fluorescence was monitored by live imaging of the apical surface throughout infection.

At 1 dpi, GFP signal was readily detectable in apically inoculated HNO-ALIs but not in basolaterally inoculated HNO-ALIs (Figure 4A-B). GFP signal steadily increased from 3 to 8 dpi in apically infected HNO-ALI cultures. In contrast, basolateral RSV inoculation resulted in minimal GFP signal until 5 to 8 dpi, where GFP signal increased in infected HNO-ALIs (Figure 4B). This pattern of minimal GFP signal until 5 to 8 dpi with basolateral inoculation, matched the kinetics data (Figure 1) where apically released virus was not observed until 5 to 8 dpi, while apical inoculation resulted in GFP signal and viral release at 1 to 2 dpi. These findings demonstrated that basolateral RSV infection results in delayed replication, spread, and viral release compared to apical infection but ultimately by 8 dpi both routes of infection result in the same infection phenotype.

**Figure 4:**
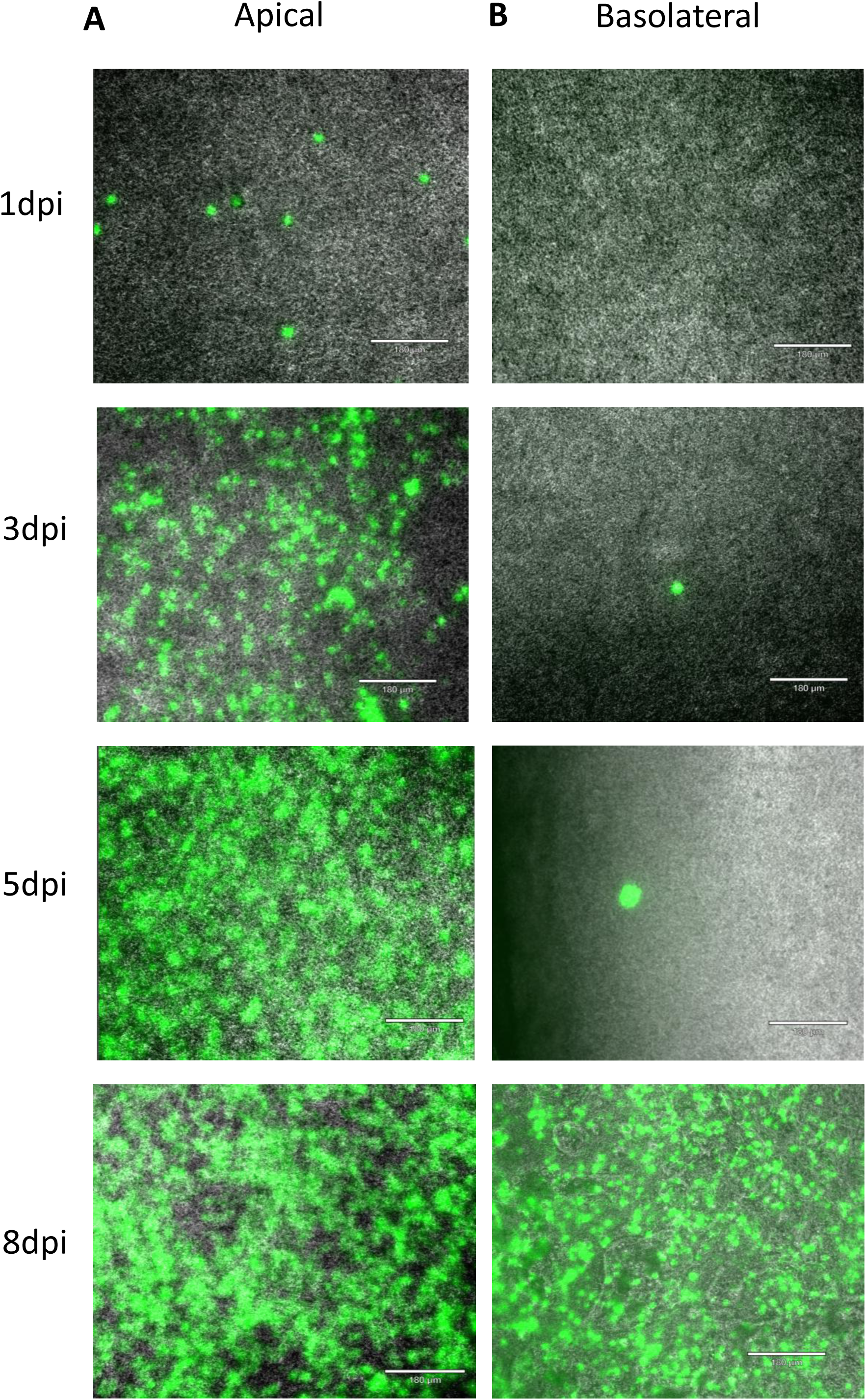
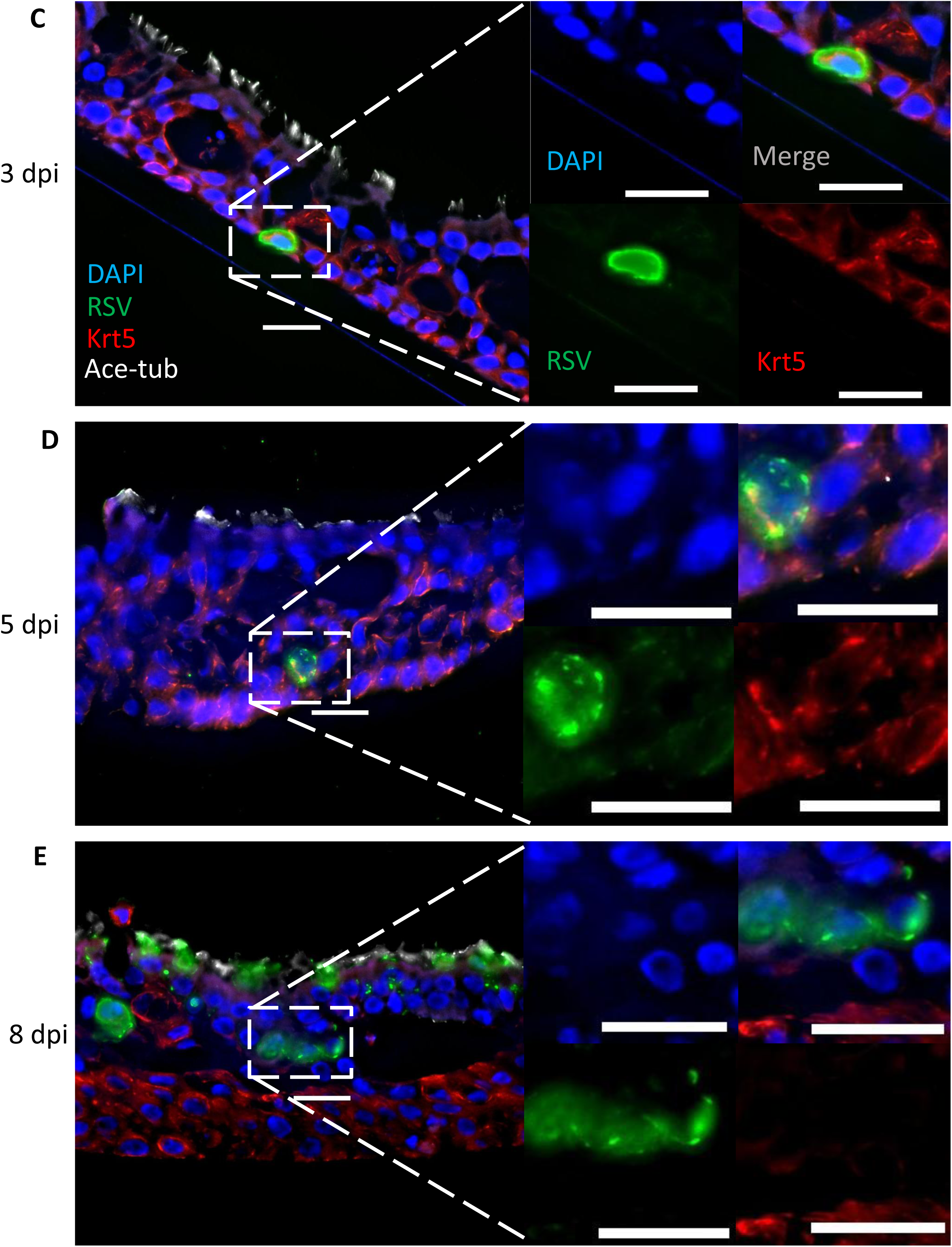
Apical and basolateral inoculation with GFP-rRSV/A2 in HNO-ALI cultures. Live microscopy images of apical HNO-ALI surface with (A) apical or (B) basolateral GFP-rRSV/A2 inoculation at 1, 3, 5, and 8 dpi from a single representative adult HNO-ALI line. Scale bar is 180 µm. (C) Representative IF images of basolateral inoculation of HNO-ALIs with GFP-rRSV/A2 at (C) 3, (D) 5, and (E) 8 dpi from a single representative adult HNO-ALI line. Basal cells – Krt5 (red), GFP-rRSV/A2 (green), Ciliated cells – Acetylated alpha-tubulin ‘Ace-tub’ (white), and nuclei – DAPI (blue). Scale bar is 20 µm.

We then hypothesized that the minimal early detection of GFP with basolateral inoculation could be the result of few initially infected basal cells before spreading to the apical ciliated cells, where GFP signal rapidly increased. We screened cross-sections of HNO-ALIs basolaterally inoculated with GFP-rRSV/A2 to investigate the spread of RSV from basal to ciliated cells. IF staining was performed on cross-sections of HNO-ALIs at 0.25, 1, 2, 3, 4, 5, and 8 dpi to identify GFP-rRSV/A2 infected basal cells. GFP-rRSV/A2 was not detected in sectioned HNO-ALIs prior to 3 dpi (data not shown). At 3, 5, and 8 dpi rare GFP-positive cells co-staining with the basal cell marker Krt5 were observed in the cross-sections (Figure 4C-E). These GFP-rRSV/A2 positive basal cells were rare (1-4 observations/4 HNO-ALI cross-sections), suggesting that RSV susceptible basal cells may represent a more specific and less prevalent basal cell subset, not identifiable by the pan-basal cell marker Krt5 alone. Interestingly, in the 8 dpi sections (Figure 4E), there were clusters of GFP-rRSV/A2 positive cells within the basal cell layer that did not stain for Krt5. However, as expected we observed frequent GFP-rRSV/A2 positive apical ciliated cells at 8 dpi, confirming ciliated cell infection occurs despite inoculating the basolateral side of the HNO-ALI.

### Basal cells are susceptible to RSV infection

To better understand basal cell susceptibility to RSV infection, we transitioned to using undifferentiated HNO-ALIs, which are comprised almost entirely of basal stem cells (18). Undifferentiated HNO-ALIs were apically or basolaterally inoculated with GFP-rRSV/A2 and GFP signal was monitored throughout infection via live imaging of the apical HNO-ALI surface. GFP-rRSV/A2 was first detectable in undifferentiated HNO-ALIs at 1 dpi and was observed through 8 dpi, regardless of inoculation route (Figure 5A-B), demonstrating basal cells are susceptible to RSV in undifferentiated HNO-ALIs. GFP-rRSV/A2 was detected early at 1 dpi in undifferentiated HNO-ALIs compared to 5 to 8 dpi in fully differentiated HNO-ALIs with basolateral inoculation, suggesting a larger population of susceptible basal cells are present in undifferentiated HNO-ALIs than in fully differentiated HNO-ALIs. However, the levels of GFP signal never increased further past 8 dpi (data not shown), indicating that basal cells are susceptible but do not support the same level of replication as the apical ciliated cells. Interestingly, there appeared to be more GFP-rRSV/A2 positive basal cells present in basolaterally inoculated undifferentiated HNO-ALIs compared to apically inoculated. This is likely due to the difference in inoculums used on the basolateral versus apical route of inoculation with an MOI of 1.0 versus 0.01, respectively.

**Figure 5:**
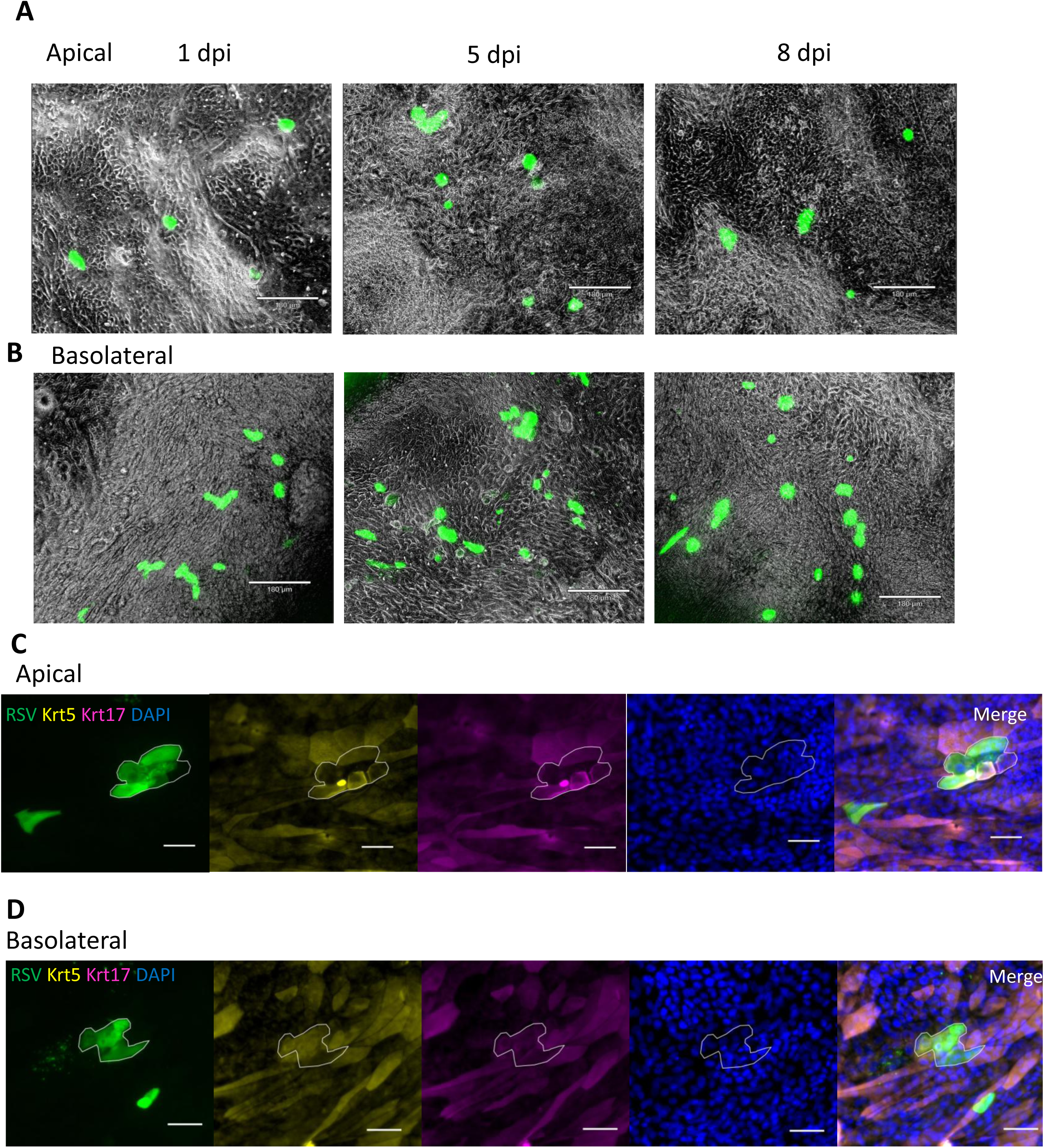
GFP-rRSV/A2 infection of basal cells in undifferentiated HNO-ALI cultures. Live microscopy images of undifferentiated HNO-ALI apical surface with (A) apical or (B) basolateral GFP-rRSV/A2 infection at 1, 5, and 8 dpi from a single representative adult HNO-ALI line. Scale bar is 180 µm. IF images of (C) apical or (D) basolateral GFP-rRSV/A2 infected undifferentiated HNO-ALI cultures at 1 dpi from a single representative adult HNO-ALI line. RSV (green), Basal cells – Krt5 (yellow) and Krt17 (pink), nuclei – DAPI (blue). Scale bar is 50 µm.

Undifferentiated HNO-ALIs expressed Krt5 and Krt17 – two canonical pan-basal cell markers (19) (Figure 5C-D, Supplemental Figure 7A). Additionally, flow cytometry confirmed that 99% of undifferentiated HNO-ALI cells co-expressed Krt5 and Krt17 (Supplemental Figure 7B). Using IF staining for Krt5 and Krt17, we determined that GFP-rRSV/A2 infected a portion of Krt5+Krt17+ basal cells in both apically and basolaterally infected HNO-ALIs (Figure 5C-D). However, given that Krt5 and Krt17 are both widely expressed basal cell markers, we expected a greater degree of RSV infection but only observed RSV in small clusters across the entire undifferentiated HNO-ALI surface. This finding further supported the concept that there is a rare subset of basal cells susceptible to RSV.

### Krt23+ activated basal cells are susceptible to RSV infection

To identify the basal cell subset infected by basolateral RSV inoculation, we revisited our previously published single-cell RNA sequencing (sc-RNAseq) dataset (20). We reported that activated basal cells (ABCs) – characterized by enriched Krt23 expression, were susceptible to low levels of RSV infection with apical inoculation of HNO-ALIs (20). Thus, we hypothesized that Krt23+ ABCs could be the susceptible basal cell population that is initially infected during basolateral RSV infection. To evaluate this, we apically or basolaterally inoculated undifferentiated HNO-ALIs with GFP-rRSV/A2 and used IF microscopy to identify GFP-rRSV/A2 infected Krt23+ ABCs.

Undifferentiated HNO-ALIs expressed both Krt23 and pan-basal marker Krt17 (Figure 6A). Krt17 was widely expressed, while Krt23 expression was restricted to a subset of basal cells. Most Krt23+ cells co-expressed Krt17+, although rare Krt23+Krt17-cells were observed (Figure 6A, white asterisks). Next, we looked for GFP-rRSV/A2 infected Krt23+ ABCs at 1 dpi. In both the apical and basolaterally inoculated undifferentiated HNO-ALIs, we observed GFP-rRSV/A2 in Krt23+ cells (Figure 6B-C). Interestingly, GFP-rRSV/A2 seemed exclusively restricted to the Krt23+ cells, with the majority of Krt17+ cells remaining uninfected. Mirroring the findings in Figure 5 where only small clusters of GFP-rRSV/A2 infected Krt5+Krt17+ cells were observed.

**Figure 6:**
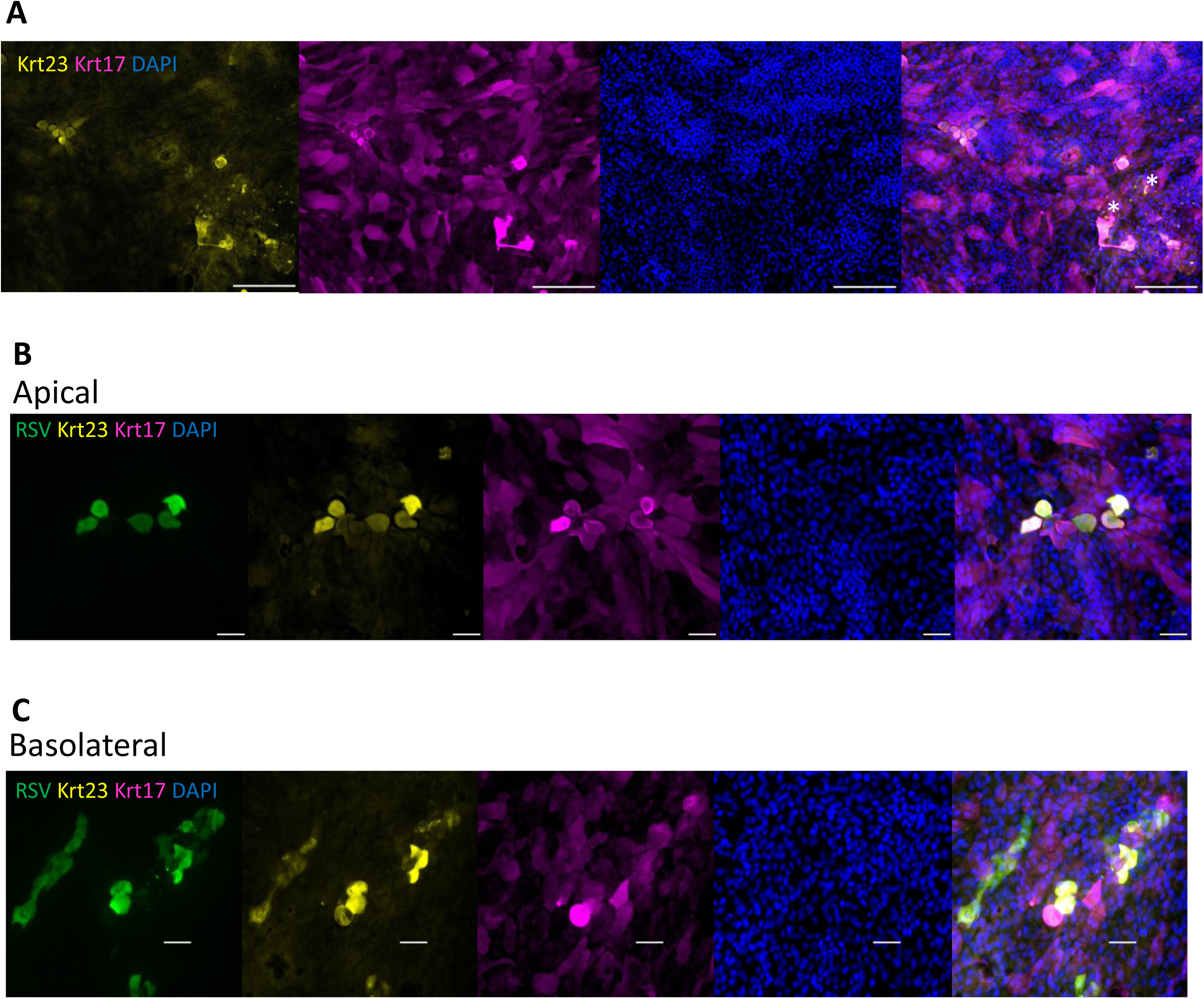
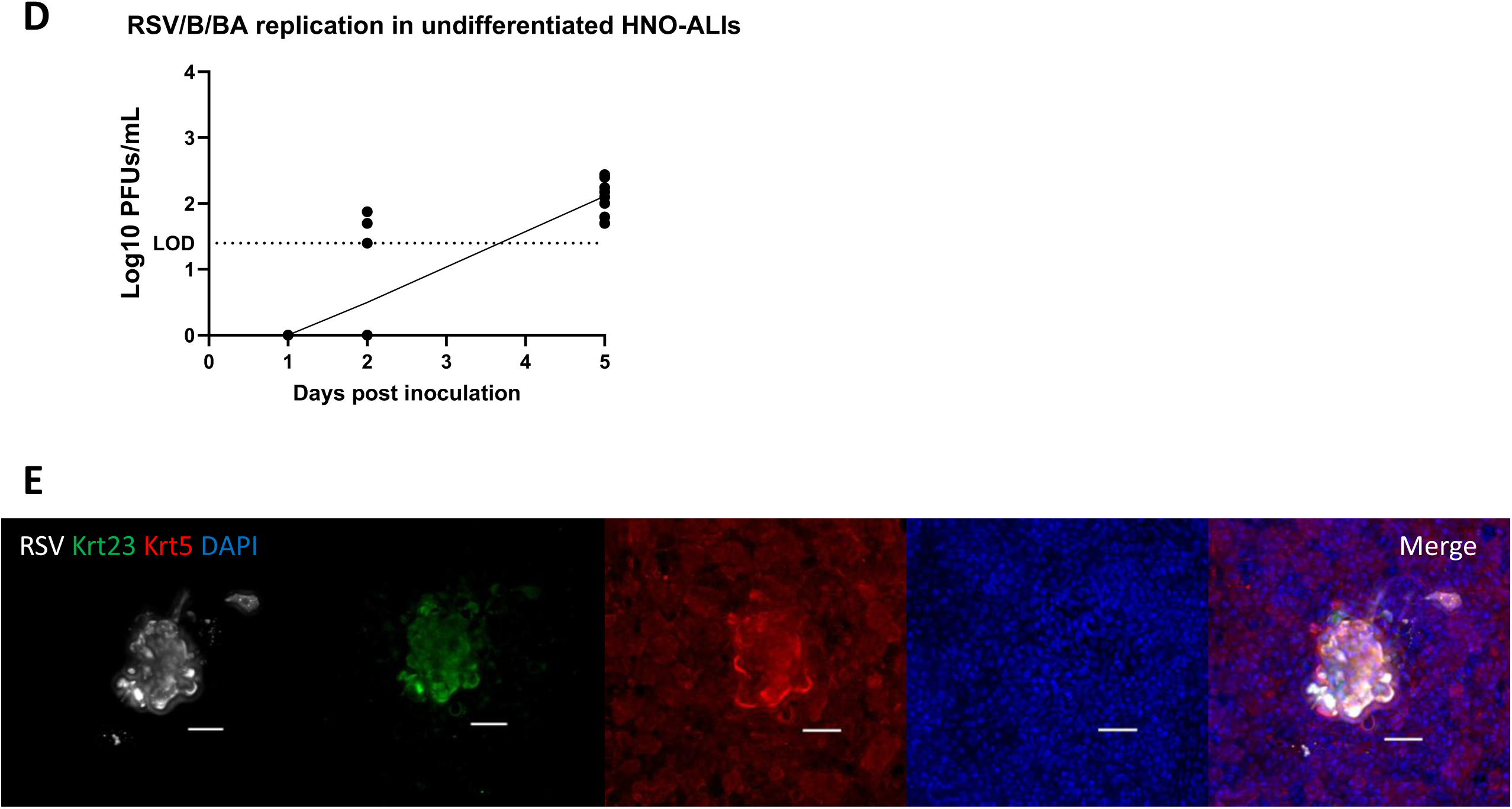
GFP-rRSV/A2 and RSV/B/BA infection of Krt23+ basal cells in undifferentiated HNO-ALI cultures. (A) Representative IF images of Krt23 expression in undifferentiated HNO-ALIs from a single representative adult HNO-ALI line. Asterisks denote Krt23+Krt17-cells. Scale bar is 180 µm. (B) Apical, or (C) basolateral GFP-rRSV/A2 inoculation in undifferentiated adult HNO-ALI cultures at 1 dpi from a single representative adult HNO-ALI line. RSV (green), basal cells – Krt23 (yellow), Krt17 (pink), nuclei – DAPI (blue). (D) Apically released infectious RSV/B/BA in undifferentiated HNO-ALIs from two adult HNO-ALI lines (4-5 technical replicates/line). LOD = limit of detection. (E) Representative IF images of RSV/B/BA infected undifferentiated HNO-ALIs from a single representative adult HNO-ALI line. Scale bar is 50 µm.

To validate this finding, we basolaterally inoculated undifferentiated HNO-ALIs with contemporaneous RSV/B/BA. At 5 dpi we detected low levels of infectious RSV/B/BA on the apical surface of undifferentiated HNO-ALIs (Figure 6D). Additionally, IF microscopy revealed that RSV/B/BA was localized to Krt23+ cells (Figure 6E). Thus, these findings demonstrate that Krt23+ ABCs are uniquely susceptible to RSV and support low levels of viral replication.

### Basolateral inoculation results in infection of Krt23+ ABCs and ciliated cells

To validate if Krt23+ ABCs were indeed susceptible to basolateral RSV infection, we transitioned back to the fully differentiated HNO-ALI basolateral infection model. 21-day differentiated HNO-ALIs of one adult HNO-ALI line were basolaterally inoculated with contemporaneous RSV/B/BA (to maximize chances of infection) and stained for flow cytometry to identify RSV infected Krt23+ ABCs and ciliated cells. Plaque assay data of apical wash samples from basolaterally inoculated HNO-ALIs showed that at 5 dpi, RSV/B/BA had infected 5/7 transwells and at 8 dpi, 7/7 transwells (Figure 7A). HNO-ALI transwells fixed at 5 and 8 dpi were stained for Krt5, Krt23, RSV F protein, and acetylated alpha-tubulin (Ace-tub) and analyzed by multi-spectral flow cytometry. The distribution of Krt23+Krt5-, Krt23-Krt5+, Krt23+Krt5+, Ace-tub+, and all other live-unstained cell types (referred to as “Other”) at 5 and 8 dpi from each transwell replicate is shown in Figure 7B.

**Figure 7:**
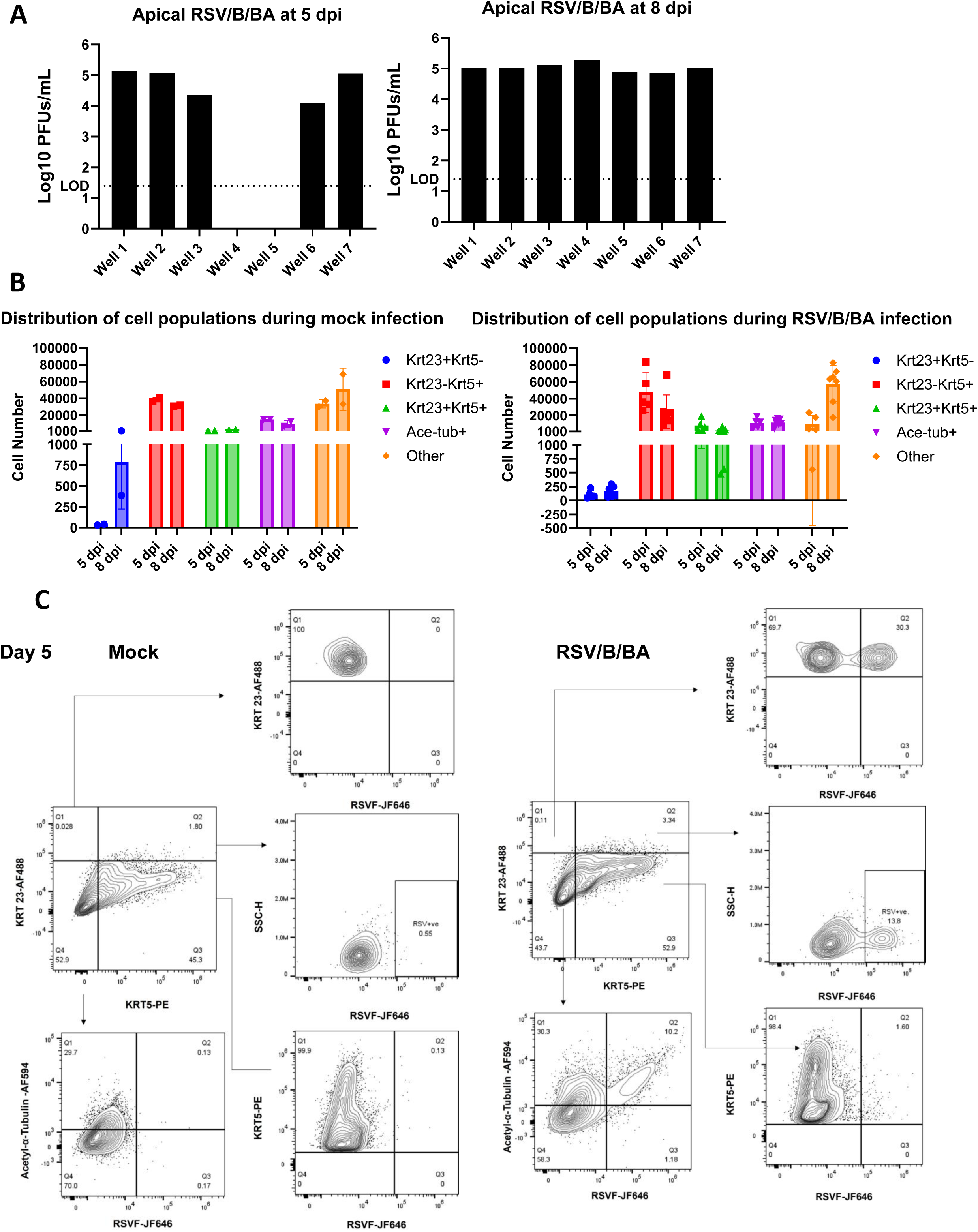
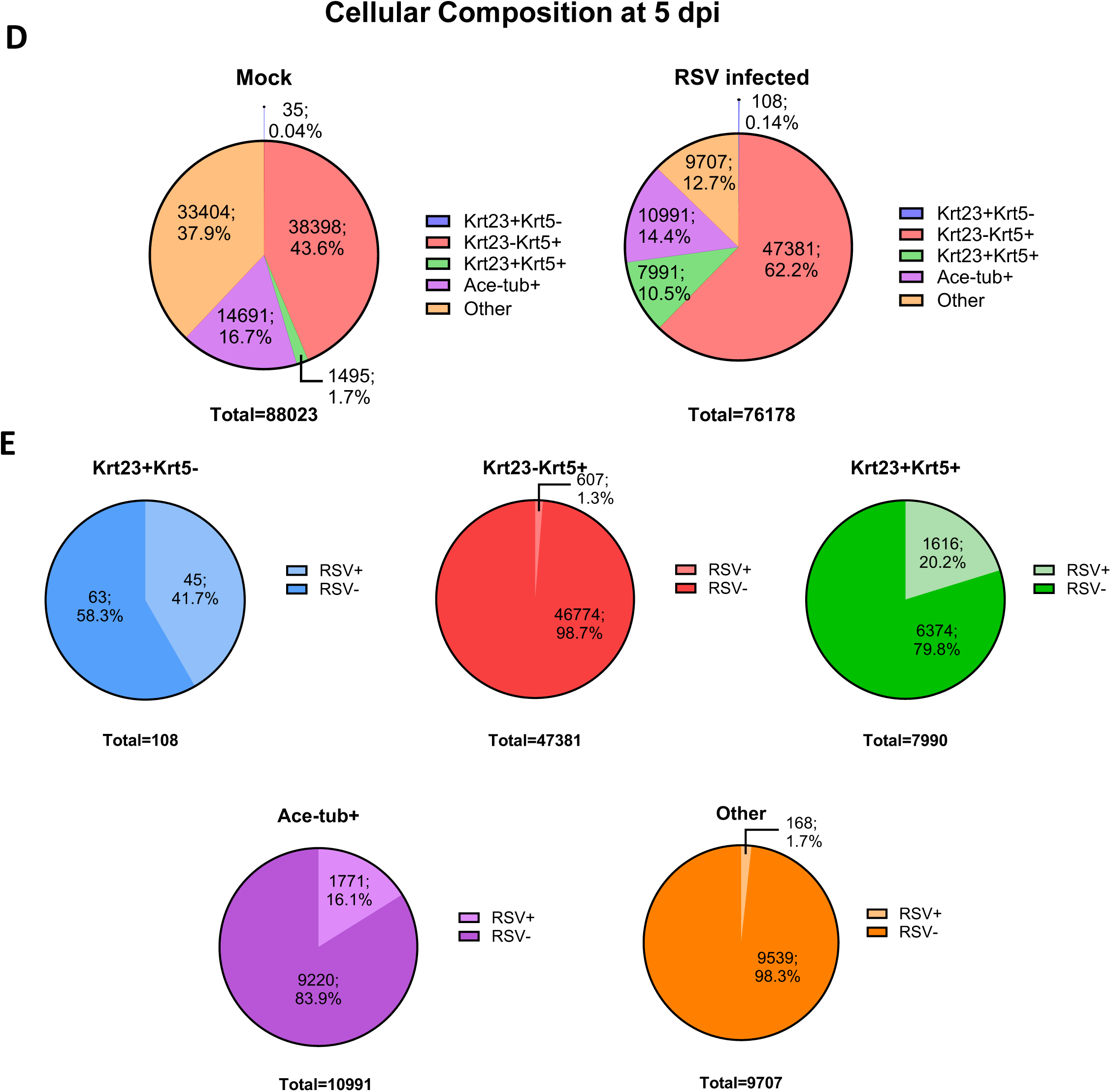
RSV selectively infects Krt23+ basal cells with basolateral inoculation of HNO-ALIs. (A) Apically released plaque forming units of RSV/B/BA from each transwell replicate of one adult HNO-ALI line at 5 and 8 dpi. LOD = limit of detection. (B) Distribution of cell populations from basolateral mock and RSV/B/BA inoculation at 5 and 8 dpi. (C) Representative spectral flow cytometry plots showing gating strategy to identify Krt5+, Krt23+, and Ace-tub+ cell populations in basolateral mock and RSV/B/BA inoculated HNO-ALIs at 5 dpi. Further gating of RSV F protein was used to determine RSV infection in each cell population. (D) Pie charts summarizing the average proportion of each cell population in basolateral mock and RSV/B/BA inoculated HNO-ALIs at 5 dpi. (E) Pie charts summarizing RSV/B/BA infection in each cell population at 5 dpi.

To determine the RSV/B/BA infected cell populations, we gated for the RSV F protein in the Krt23+Krt5-, Krt23-Krt5+, Krt23+Krt5+, and Ace-tub+ populations at 5 and 8 dpi (Figure 7C, Supplemental Figure 8A). There was no evidence of RSV infection in any of the cell populations in the basolateral mock infected HNO-ALI transwells at 5 and 8 dpi (Figure 7C, Supplemental Figure 8A). At both 5 and 8 dpi, we observed RSV infection predominantly in the Krt23+Krt5-, Krt23+Krt5+, and Ace-tub+ populations (Figure 7C, Supplemental Figure 8A). Thus, demonstrating that basolateral inoculation of HNO-ALIs leads to infection of Krt23+ ABCs and ciliated cells. Data from RSV/B/BA infected transwells at each timepoint were pooled and averaged for all subsequent analysis.

At 5 dpi, Krt23⁺Krt5⁻ cells represented the smallest population, comprising <1% of total cells (0.03–0.04% in mock and 0.03–0.27% in RSV/B/BA infected HNO-ALIs), whereas Krt23⁻Krt5⁺ cells accounted for the largest population, averaging 43.6% (43.18–44.12%) in mock and 62.2% (40.26–72.64%) in RSV/B/BA infected HNO-ALIs (Figure 7D). Krt23⁺Krt5⁺ cells accounted for a small fraction of total cells, averaging 1.7% (1.63–1.78%) in mock and 10.5% (3.34–22.96%) in RSV/B/BA infected HNO-ALIs (Figure 7D). Ace-tub⁺ ciliated cells represented 16.7% (15.60–17.92%) of mock-infected cells and 14.4% (7.04–19.84%) of RSV/B/BA infected HNO-ALIs. All other live unstained cells (“Other”) accounted for 37.9% (36.13–39.58%) in mock and 12.7% (0.67–36.23%) in RSV/B/BA infected HNO-ALIs (Figure 7D). Similar distributions were observed at 8 dpi, although a larger proportion of “Other” cells and a smaller proportion of Krt5⁺ cells were detected in both mock and RSV/B/BA infected HNO-ALIs, compared to 5 dpi (Supplemental Figure 8B). Possibly due to an RSV induced increase in goblet cells (12, 13).

We next assessed the frequency of RSV infection within each defined cell population at 5 dpi. Interestingly, despite Krt23+Krt5-cells accounting for the smallest percentage of total cells, they were the most permissive to RSV/B/BA with an average of 41.7% (7.69-69.16%) of Krt23+Krt5-cells being RSV+ (Figure 7E). In contrast, Krt23-Krt5+ cells are the majority cell type but were the least permissive to RSV/B/BA with an average of 1.3% (0.34-1.77%) of Krt23-Krt5+ cells being RSV+ (Figure 7E). Dual positive Krt23+Krt5+ cells were more permissive than Krt23-Krt5+ alone, with an average of 20.2% (3.62-28.32%) of cells being RSV+, indicating the presence of Krt23 could be a marker of increased susceptibility to RSV (Figure 7E). As expected, we also detected infected ciliated cells, with 16.1% (3.82-39.56%) of Ace-tub+ cells being RSV+. Lastly, there was a very small population of 1.7% (0.71-23.26%) of “Other” cell types that were RSV+. Similar results were seen in the 8 dpi transwells (Supplemental Figure 8C) where Krt23+Krt5-, Krt23+Krt5+, and Ace-tub+ cells accounted for the most RSV infected populations. Taken together, these results show that Krt23+ ABCs are highly susceptible to RSV infection while Krt23-Krt5+ basal cells are resistant to infection, and that basolateral inoculation can result in infection of the rare Krt23+ ABCs with subsequent spread of the virus to the apical ciliated cells.

## Discussion

RSV is responsible for millions of annual LRTIs globally. RSV is estimated to affect the lower airways in 10-45% of children infected with RSV <5 years of age, resulting in bronchiolitis and pneumonia (21–23). In hematopoietic stem cell transplant (HSCT) patients, approximately 40-60% of RSV infections will progress to LRTIs, with an estimated mortality rate of 80% in those patients (24). Additionally, in older adults, the clinical manifestation of LRTI symptoms were reported in 30-90% of RSV cases (25, 26) Therefore, there is a crucial need to further understand the routes of RSV infection and potential mechanisms of spread to the lower respiratory tract to fully understand the pathogenesis of the virus. We thus sought to determine if RSV was able to infect the respiratory epithelium via the basolateral route of infection in an *ex vivo* human nose organoid model.

We found that contemporaneous RSV/A/ON and RSV/B/BA strains infect both adult and infant-derived HNO-ALIs with basolateral inoculation, but at different frequencies. Notably, basolateral infection required at least one-hundred-fold higher inoculum compared to apical infection (1.0 vs 0.01 respectively). A higher inoculum is potentially needed for basolateral infection to increase the odds of RSV encountering rare Krt23+ ABCs to initiate infection. Basolateral RSV infection resulted in a prolonged eclipse phase with no detectable RSV shedding into the apical lumen for approximately 5 days, followed by high levels of apically released infectious virus from infected ciliated cells between 5 to 8 dpi. Importantly, basolateral infection followed by apical release of RSV did not occur in all transwells, even though RSV/B/BA infected at a significantly higher frequency compared to RSV/A/ON. The observed variation in transwell-to-transwell infection outcome, even within the same donor line, suggests that basolateral infection depends on the relative abundance or activation state of the rare Krt23⁺ basal cell subpopulation.

Our findings contrast with those of *Zhang et al*. in 2002, who reported that RSV could not infect human airway epithelial cells via basolateral inoculation (16). However, in their study, basolateral infection outcome was only determined at 24 hours post infection, whereas our data demonstrate it can take up to 8 days for detection and apical release of the virus to occur. Similar findings were observed by *Zhang et al*. in 2005, where productive basolateral infection with parainfluenza virus was not seen in differentiated human airway epithelial cells 48 hours post infection (27). However, they observed that basal cells of undifferentiated cultures were susceptible to parainfluenza virus infection (27). These findings align with our observation that a prolong eclipse time is needed for apical detection of RSV, and that direct inoculation of basal cells results in productive infection.

A novel finding of our study was the observed difference in infectivity between RSV/A/ON and RSV/B/BA following basolateral inoculation. The two viral strains used in our studies are contemporaneous RSV strains from the two dominant, circulating ON and BA genotypes, which are both characterized by a duplication in the distal third of the attachment (G) glycoprotein (28). We found that with basolateral inoculation of adult and infant-derived HNO-ALIs, RSV/B/BA infected approximately 80% of the transwells, while RSV/A/ON infected approximately 25%. Additionally, this difference in infectivity was reproducible with different strains of the ON and BA genotypes. Notably, this is the first demonstration of a significant difference in infection outcome between contemporaneous RSV strains from the two subgroups. This difference was observed only with basolateral inoculation and was not observed in our prior studies using apical infection of HNO-ALIs (12, 13, 28). To date, most studies have utilized prototypic strains of both RSV/A and RSV/B or have compared strain variability within the same RSV subgroup (29–31). Our study underscores the importance of utilizing contemporaneous RSV isolates of both subgroups for *in vitro* comparison studies to fully understand RSV pathogenesis.

The two RSV subgroups are antigenically distinct from one another, with the most heterogeneity occurring in the G protein, which aids viral binding to an attachment receptor (32). We hypothesize that the difference in basolateral infection outcome may be due to differences in the G protein that grant RSV/B/BA an advantage over RSV/A/ON. It has been shown that airway basal cells express heparan sulfated proteoglycans (HSPGs), a known attachment receptor that binds the G-protein via the heparin binding domain (33, 34). Therefore, it is possible that RSV utilizes HSPGs as an attachment receptor for basal cell infection, but RSV/B/BA has an intrinsic advantage over RSV/A/ON for receptor binding in this infection context. Additionally, our prior single-cell RNA sequencing study identified transcripts of other known RSV co-receptors including intercellular adhesion molecule-1 (ICAM-1), insulin-like growth factor 1 receptor (IGF1R), and HSPGs that were expressed in basal cells that may play a role in infection ((20). Specifically, we saw high levels of ICAM-1 expression on Krt23+ ABCs. In addition, RSV/B/BA may be more effective in overcoming intrinsic antiviral defenses of basal cells compared to RSV/A/ON. To this end, future studies are needed to investigate which RSV attachment and entry receptors are expressed on basal cells and how differential receptor usage may explain the observed subgroup differences in basolateral infection outcome.

The apical ciliated cells of the respiratory epithelium are the predominant target cell population of RSV. To ensure that infection of the ciliated cells with apical release of the virus was not occurring by translocation of the virus through loss of tight junction integrity, we monitored TEER throughout infection. TEER measures epithelial cell integrity by assessing how much electrical resistance a cell monolayer provides, with higher TEER values reflecting stronger barriers and lower values suggesting compromised barriers. With both apical and basolateral inoculation, we saw that TEER increased with RSV infection, but to a greater degree with apical infection. This mirrors the findings published by *Kast et al*. who reported that RSV infected human bronchial epithelial cells had increased TEER and decreased paracellular flux throughout infection, suggesting an increase in epithelial barrier integrity as an early epithelial protective response to infection (35). However, another study demonstrated that in mice, RSV decreased TEER, increased epithelial permeability, and altered the expression of several tight junction proteins (36). This difference in findings is most likely due to the lack of a cellular immune component and a limitation of epithelium-only based studies, whereas in animal studies, the RSV induced immunopathology is a key contributor to altered epithelial barrier integrity (36). In fact, *Deng et al*. observed that with the addition of neutrophils to RSV infected airway epithelial cells, TEER and barrier function decreased compared to RSV infection alone, where barrier function was maintained (37). In our study, we did not observe a loss of epithelial integrity with basolateral inoculation, indicating that infection of the apical ciliated cells was not due to a loss of barrier function. Instead, infection was most likely due to viral infection of a rare basal cell type with subsequent intracellular spread to the apical ciliated cells.

We identified a rare basal cell type, Krt23+ ABCs, as the predominant basal cell type susceptible to basolateral RSV infection. Studies in the preterm lamb and infant baboon model have reported occasional RSV infected basal cells in the airway epithelium, that were not further characterized (38, 39). RSV infection of basal cells has remained largely unstudied until *Persson et al*. demonstrated that basal cells are susceptible to RSV in a scratch injury model, and that infection of basal cells can skew the differentiation of the epithelium to favor a secretory phenotype (17). Our study differs from that of *Persson et al*. in that we did not damage the epithelium to expose the basal cell layer to RSV by apical inoculation. Rather, we exposed the basal cell layer of intact HNO-ALIs to RSV by direct basolateral inoculation and yet were able to see productive infection. Additionally, *Persson et al*. reported p63+ (a pan-basal cell marker) basal cells, were susceptible to RSV, while we were able to refine this observation by identifying that Krt23+ ABCs are the predominant basal cell subset susceptible to RSV infection.

Using undifferentiated HNO-ALIs we observed that Krt23+ ABCs are susceptible to contemporaneous RSV/B/BA but do not support high levels of viral replication. Therefore, Krt23+ ABCs likely allow for limited viral replication to initiate infection, but the spread to ciliated cells is necessary to sustain the infection and allow robust viral replication. Additionally, we observed a gradient in RSV infectivity with the highest infection rate in Krt23+Krt5-ABCs followed by Krt23+Krt5+ basal cells while Krt23-Krt5+ basal cells were resistant to infection. Indicating that RSV susceptibility differs across different basal cell subpopulations.

Krt23+ ABCs (also referred to as para/supra basal cells) remain an understudied basal cell population in human airways. Krt23+ ABCs are described in literature as being more differentiated than Krt5+ basal cells and serving as an early progenitor cell type, residing between the basal and luminal cell layers (19, 40). Krt23 is a proliferation marker and has been shown to be specifically expressed in parabasal cells (19, 41, 42). *Boers et al*. reported that parabasal cells make up 3-7% of epithelial cells/millimeter of upper airway, while Krt5+ basal cells account for approximately 30% of cells (43). Our findings are consistent with these data, in that Krt23+ ABCs comprised a very small percentage of basal cells while Krt5+ cells were the dominant basal cell type. Therefore, our data aligns with previous studies that suggest Krt23+ ABCs are a rare cell type found in the airway epithelium. However, this cell population and what specifically renders them more susceptible to RSV compared to other basal cell subsets remain unclear. It is possible that due to their proliferative nature, Krt23+ basal cells are more susceptible to RSV compared to the quiescent basal cells, allowing for viral propagation. Alternatively, Krt23+ is a unique basal cell marker that could accompany the presence of an RSV entry receptor, allowing Krt23+ cells to be susceptible to RSV infection. We know from our prior single-cell RNA sequencing study along with studies from other groups, that several RSV entry receptors are expressed on basal cells (20, 27, 33, 44). Further studies will be needed to identify the specific receptor utilized by RSV for infection of Krt23+ ABCs and to determine how receptor expression patterns differ across basal cell subsets.

Despite the clinical relevance and consistency of our findings, there are several limitations to the HNO-ALI model. Firstly, the HNO-ALI model only recapitulates the epithelium of the upper airways. Validating these findings in organoid models of the lower airways will be important to understand the lower airways’ susceptibility to basolateral RSV infection and the basal cell susceptibility. Secondly, our current HNO-ALI model lacks immune cells, submucosa, and endothelial structures, which limit our ability to fully mimic the basolateral milieu of the airway epithelium *in vivo*. In the future, more complex organoid model systems will be needed to better model the basolateral route of infection. However, despite these limitations, we consistently observed that the basolateral route of exposure is a viable infection route for RSV infection and due to the cellular heterogeneity found in HNO-ALIs, we identified Krt23+ ABCs to be a uniquely susceptible basal cell subset to RSV.

In summary, we determined that the respiratory epithelium is susceptible to the basolateral route of infection by RSV with an unusually prolong eclipse phase, there are differences in infection outcome between contemporaneous strains of the RSV subgroups suggesting subtle differences in viral attachment and cell entry, and we identified a new RSV susceptible cell population – Krt23+ ABCs. These findings suggest that RSV could utilize an alternate route of infection to contribute to LRTI by infecting a unique basal cell population within the basal cell layer of the distal airways. Investigating alternate routes of infection is critical to understanding the susceptibility of the different cell populations of the airway epithelium and their contribution to the RSV pathogenesis. Thus, expanding our understanding of how different routes of infection contribute to disease progression, viral tropism, host immune response, and long-term sequelae.

## Materials and Methods

### HNO-ALI Cell Lines

Eight adult lines (HNO-ALI 204, 02, 918, 919, 929, 934, 923, 930) and eight infant lines (HNO-ALI 9007, 9009, 9003, 9005, 9002, 9004, 9006, 9008) were used throughout this study. HNO-ALI cell lines were generated as previously described (12, 13). Briefly, nasal wash and mid-turbinate (M-T) swab samples were collected from adults and infant participants – after obtaining signed, informed consent by an adult or legal guardian through an approved protocol by the Baylor College of Medicine Institutional Review Board. Nasal wash combined with M-T swab samples were placed in digestion media (10 mL airway organoid (AO) medium + 10 mg Collagenase (Sigma C9407) +100 µL Amphotericin B), strained to remove debris, and washed. Cells were pelleted down and resuspended in Matrigel^®^ (Corning, NY) and plated as 3D HNOs in growth media. 3D HNOs were expanded for 3-4 days (infant) and 5-7 days (adults). After 4 or more rounds of passage, 3D HNOs were enzymatically and mechanically disrupted to make a single cell suspension and were seeded onto transwells^®^ (Corning, NY) at a density of 1-3 x 10^5^ cells/well. Cell monolayers were maintained in AO propagation media with endothelial growth factor (EGF, Peprotech-AF-100-15) containing 10 µM Y-27632. After 4 days in a liquid environment, the monolayers were transferred to an air-liquid interface (ALI) environment with AO differentiation (AO Diff) media (PneumaCult-ALI, STEMCELL Technologies) in the basolateral compartment of the transwell. HNO-ALI cultures were maintained in a humidified incubator at 36 °C with 5% CO_2_. AO Diff media was changed every 4-5 days, and the cultures were maintained for 21 days to form a stratified multi-cellular ciliated epithelium. For undifferentiated HNO-ALI cultures, the monolayers were grown 2-4 days on transwells in AO propagation media and switched to ALI conditions and AO Diff media at the time of infection.

### Viruses

Working pools were generated of RSV/A/USA/BCM813013/2013 (genotype ON), HRSV/A/USA/TX-HOU-R06687/2021 (genotype ON), RSV/B/USA/BCM80171/2010 (genotype BA), and HRSV/B/USA/TX-HOU-IP0035B/2018 (genotype BA). Viruses referred to as RSV/A/ON, RSV/A/ON-Hou, RSV/B/BA, and RSV/B/BA-IP, respectively. All viruses were passaged twice in human epidermoid carcinoma larynx cells (HEp-2, ATCC) to generate working pools. GFP-rRSV/A2 was generated as previously described and generously gifted by Dr. Ursula Buccholz (45). Briefly, recombinant RSV derived from the WT RSV/A2 strain, expressing GFP between the P and M genes was passaged twice in HEp-2 cells to create working pools of GFP-rRSV/A2. Sucrose purified GFP-rRSV/A2 was purified by ultracentrifugation on a 30%-60% sucrose gradient as previously published (46).

### Viral Infection and Sample Collection

Basolateral Infection: 21-day fully differentiated HNO-ALI cultures were basolaterally inoculated with 600 µL of either RSV/A/ON, RSV/B/BA, or GFP-rRSV/A2 diluted in AO Diff media at a multiplicity of infection (MOI) of 1.0 based on total cell count or mock inoculated with AO Diff media. The inoculum was incubated in the basolateral compartment for 1.5 hours at 36 °C with 5% CO_2_ and then removed and replaced with 600 µl of AO Diff media. 2-3 technical replicates of HNO-ALI transwells were used for measuring viral kinetics at 0.25, 1, 2, 5, and 8 days post-inoculation (dpi). At each timepoint, apical wash samples were collected with three consecutive 200 µL washes of AO Diff media. The total 600 µL wash volume was mixed 1:1 with 15% glycerol/Iscoves media, aliquoted, snap-frozen in a dry ice alcohol bath, and stored at –80 °C. Basolateral samples were collected by removing the 600 µL basolateral media, mixing 1:1 with 15% glycerol/Iscoves media, aliquoted, snap-frozen in a dry ice alcohol bath, and stored at –80 °C, and the epithelial cell layer was fixed with 4% PFA for immunofluorescent microscopy and histology. Alternatively, at 0.25, 1, 2, 3, 4, 5, 6, and 8 dpi the epithelial cell layer was enzymatically removed with 200 µL trypsin (Corning) and mixed with 400 µL viral lysis buffer (Revvity) for detection of intracellular viral RNA. All HNO-ALI infections were performed independently on different days/times.

Apical Infection: Fully differentiated HNO-ALI cultures were apically infected with 30 µl of a 0.01 MOI of either RSV/A/ON, RSV/B/BA, or GFP-rRSV/A2 diluted in AO Diff media or mock inoculated with AO Diff media for 1.5 hours at 36 °C with 5% CO_2_ and then the inoculum was removed followed by the sample collection protocol described above.

### PCR and Plaque Assays

Viral RNA was extracted using either a mini viral RNA kit (Qiagen Sciences) on an automated QIAcube platform or Chemagic Viral RNA 300 Kit H96 kit on a Chemagic^TM^ 360 automated platform according to the manufacturer’s instructions (47). Viral RNA was detected and quantified using real-time polymerase chain reaction with primers targeting the nucleocapsid (N) gene of RSV (47). Infectious RSV (PFUs/mL) was measured using a quantitative plaque assay as previously described (48). The lower limit of detection was 25 PFUs/mL. Samples that were below the limit of detection were assigned the value of 1 PFUs/mL.

### TEER measurements

TEER was measured with the EVOM3 Volt/Ohm meter (World Precision Instruments). Baseline TEER prior to infection was taken from 21-day fully differentiated HNO-ALI cultures. HNO-ALI cultures of two adults (918, 204) and two infants (9007, 9009) were mock, basolateral, or apically infected with RSV/A/ON or RSV/B/BA as described above. Duplicate wells of each line were used for each infection condition. TEER was measured at 0.25, 1, 2, 5, and 8 dpi with duplicate measurements per well. Simultaneously, apical wash and basolateral media samples were collected as described above. RSV replication kinetics and TEER measurements were averaged for adult and infant HNO-ALI lines.

### Immunohistochemistry (IHC) and Immunofluorescence staining

HNO-ALI cultures were fixed in image-iT™ Fixative Solution (4% formaldehyde) [Thermo Fisher Catalog number: FB002] for 15 minutes followed by dehydration in ethanol series (30%, 50%, and 70%) for 30 minutes each at room temperature. The transwell membranes were then paraffin embedded and sectioned for standard hematoxylin and eosin (H&E) or immunofluorescence (IF) staining. For IF staining, sections were deparaffinized in Histo-Clear, followed by ethanol washes (100>100>90>70%). The slides were rehydrated in PBS, followed by heat-induced antigen retrieval in 10 mM sodium citrate buffer. Sections were then rinsed in water and blocked with 5% bovine serum albumin (BSA) in PBS. Sections were incubated with the following primary antibodies: keratin 5 for basal cells (1:2000, Krt5; BioLegend, Catalog number: 905503), acetylated alpha tubulin for ciliated cells (1:1000, Santa Cruz, Catalog number: sc-23950), goat polyclonal antibody specific for RSV (1:1000, Abcam, Catalog number: ab20745), or goat polyclonal antibody specific for GFP (1:1000, Abcam, Catalog number: ab290) overnight at 4 °C. Primary antibodies were washed four times in PBS for 5 minutes each and incubated with secondary antibodies: donkey anti-rabbit 568 (Invitrogen, Catalog number: A-10042), donkey anti-goat 647 (Invitrogen, catalog number: A-21447), donkey anti-goat 488 (Invitrogen, Catalog number: A-32814) and 4′,6-diamidino-2-phenylindole (DAPI) for 2 hours at room temperature. Sections were washed three times with PBS for 5 minutes each, rinsed with water, mounted in Prolong^TM^ Gold (Life Technologies, Catalog number: P36930) and cured overnight at room temperature. Slides were stored at 4 °C.

For IF staining of the whole transwell membrane, undifferentiated HNO-ALI transwell membranes were fixed and dehydrated as described above. The membranes were then rehydrated in ethanol (50%, 30%) and PBS for 30 minutes each. The membranes were then permeabilized in 0.1% Triton-X in PBS for 10 minutes with gentle agitation, followed by washing four times with tris-buffered saline with tween® 20 (TBS-T) diluted to 0.05% in distilled water. The membranes were blocked with 5% BSA in PBS for 2 hours, followed by overnight incubation at room temperature with the following primary antibodies in 5% BSA: keratin 5 (1:2000, Krt5 for basal cells; BioLegend, Catalog number: 905503), keratin 17 (1:1000, Krt17 for basal cells; Santa Cruz, Catalog number: 393002), keratin 23 (1:500, Krt23 for activated basal cells; Santa Cruz, Catalog number: 22590AF488), goat polyclonal antibody specific for GFP (1:1000, Abcam, Catalog number: ab290), or goat polyclonal antibody specific for RSV (1:1000, Abcam, Catalog number: ab20745). Primary antibodies were washed four times with TBS-T and membranes were incubated overnight at room temperature with secondary antibodies in 5% BSA: donkey anti-rabbit 568 (Invitrogen, A-10042), donkey anti-mouse 568 (invitrogen, Catalog number: A-10037), donkey anti-goat 647 (Invitrogen, Catalog number: A-21447). Secondary antibodies were washed four times with TBS-T and membranes were incubated with DAPI for 2 hours. DAPI was washed four times with TBS-T and the membranes were mounted in Prolong^TM^ Gold and cured overnight at room temperature.

### Immunofluorescent Imaging

Sectioned samples were imaged using high resolution Cytiva DVLive or Olympus IX83 epifluorescence deconvolution microscopes. Images were collected with a 40x/0.95NA objective with a 10 µm z stack (using optical sections at the recommended Nyquist for each objective). 3D images were deconvolved using a quantitative image restoration algorithm. Max intensity projections were used for image analysis and processed using Fiji (49). For H&E-stained sections, images were obtained on a Nikon CiL Brightfield microscope at 40x/0.65NA. Duplicate samples of each transwell were imaged with two images obtained per sample. Whole transwell membranes were imaged using the Revolve Fluorescent Microscope (Echo) at 10x/0.30NA and 20x/0.45NA with a 10 µm z-stack taken (using optical sections at the manufacturer’s recommended step size). Max intensity projections were used for image analysis and processed using Fiji (49).

### Undifferentiated HNO-ALI infection and imaging

GFP-RSV: Undifferentiated HNO-ALI cultures were apically or basolaterally inoculated with sucrose purified GFP-rRSV/A2 (kindly provided by Ursula J Buchholz (National Institute of Allergy and Infectious Diseases, Bethesda, MD) at an MOI of 1 or apically inoculated at an MOI of 0.01. For undifferentiated cultures, cells were switched from AO propagation media to AO Diff media and ALI conditions at the time of infection. After 1.5 hours the inoculum was removed and replaced with AO Diff media. Duplicate HNO-ALI transwells were used for each time point and GFP expression was monitored daily with the Revolve Fluorescent Microscope (Echo) at 10x/0.25. At 0.25, 1, 2, 3, 4, 5, and 8 dpi. Apical wash and basolateral media samples were collected for viral kinetics as described above and the epithelial cell layer was fixed for immunofluorescent imaging. RSV/B/BA: Undifferentiated HNO-ALI cultures were basolaterally inoculated with RSV/B/BA as described above. Apical wash samples were obtained at 1, 2, and 5 dpi for viral kinetics, and the epithelial cell layer was fixed for immunofluorescent imaging.

### Intracellular Staining and FACS Analysis

FACS analysis was performed to determine the proportions of different epithelial cell populations in HNO-ALIs and to assess RSV infection within each cell population. Briefly, 21-day fully differentiated HNO-ALI cultures from one adult line were basolaterally inoculated with RSV/B/BA as described above. Cells were then washed with PBS and gently dissociated from culture wells using TrypLE Select (Thermo Fisher Scientific, Catalog number: 12604013), then centrifuged at 600 g for 5 min at 4 °C. Cells were transferred to 96-well V-bottom plates (Thermo Fisher Scientific, Catalog number: 277143), and stained for viability using the LIVE/DEAD™ Fixable Violet Dead Cell Stain Kit (Invitrogen, Catalog number: L34955) for 10 minutes at room temperature in the dark. Cells were then washed with FACS buffer (0.5% BSA, 1 mM EDTA in PBS) and permeabilized using the Cytofix/Cytoperm™. Fixation/Permeabilization Kit (BD Biosciences, Catalog number: 554714), following the manufacturer’s instructions. Immunostaining for intracellular proteins was performed using the following primary conjugated antibodies: Krt5 (1:5000, Abcam, Catalog number: ab224985), Krt17 (1:100, Abcam, Catalog number: ab185032), Krt23 (1:100, Novus Biologicals, Catalog number: NBP2–22590AF488), acetylated α-tubulin (1:100, Santa Cruz, Catalog number: sc-23950 AF594), and RSV-F protein (1:100, Novus Biologicals, Catalog number: NB100–64495JF646). Antibodies were incubated for 30 minutes on ice. Following staining, cells were washed twice with BD Perm/Wash buffer and resuspended in FACS buffer prior to acquisition on a Cytek Aurora CS spectral flow cytometer. Data were analyzed using FlowJo 10 Software (BD Biosciences).

### Statistical Analysis

Statistical analysis was performed using Prism software v10 (Graphpad). Infection status of HNO-ALI transwells was determined at 5 and 8 dpi, in which a transwell was deemed infected by the detection of infectious virus on the apical surface. The total number of infected and uninfected wells were counted, and statistical significance was determined by Fisher’s Exact Test. To determine the statistical significance of RSV infection on TEER, a two-way ANOVA was performed with Dunnett’s correction for multiple comparisons. Statistical significance was indicated for p-values <0.05.

### Data Availability

A supporting data values file with all reported values will be included as part of the supplementary material.

## Acknowledgements

This work was supported by funds from the National Institutes of Health (NIH) grants U19 AI116497 (SEB, VA, PAP) and U19AI144297 (SEB, VA, PAP). Imaging for this project was supported by the Integrated Microscopy Core at Baylor College of Medicine and the Center for Advanced Microscopy and Image Informatics (CAMII) with funding from NIH (DK56338, CA125123, ES030285, S10OD030414), and CPRIT (RP150578, RP170719). Tissue processing services for this project were supported by the Digestive Disease Center core for Tissue Analysis and Molecular Imaging supported in part by PHS grant P30DK056338. We thank Dr. Ursula Buccholz for generating and providing GFP-rRSV/A2. Additionally, we thank Amal Kambal, Alejandra Rivera-Tostada, and Xi-Lei Zeng for their generation and maintenance of the HNO-ALI cultures and Pamela Parsons for her tissue processing services.

## Author Contributions

AM designed the project, performed experiments, analyzed data, and wrote the manuscript draft. DN performed flow cytometry, analyzed data and edited the manuscript. EMS and GA performed infections, viral kinetics measurements, and edited the manuscript. EN consented subjects, obtained nasal washes and M-T swab samples from subjects for generating HNO lines, and edited the manuscript. SEB supervised established organoid cultures and edited the manuscript. PAP and VA designed the project, analyzed data, obtained funding, and wrote and edited the manuscript. All authors reviewed and approved the final manuscript.

## Figure Legends

**Supplemental Table 1:** Demographics of adult and infant HNO-ALI donors.

| **INFANT HNO-ALIs** |  |  |
| --- | --- | --- |
| **Sample ID** | **Gender** | **Age range** |
| HNO9002 | F | 12-24 months |
| HNO9003 | F | <12 months |
| HNO9004 | F | <12 months |
| HNO9005 | M | 12-24 months |
| HNO9006 | M | 12-24 months |
| HNO9007 | F | <12 months |
| HNO9008 | M | 12-24 months |
| HNO9009 | M | 12-24 months |
| **ADULT HNO-ALIs** |  |  |
| **Sample ID** | **Gender** | **Age range** |
| HNO204 | F | 65+ years |
| HNO02 | M | 50-64 years |
| HNO918 | F | 18-45 years |
| HNO919 | F | 18-45 years |
| HNO923 | M | 18-45 years |
| HNO929 | M | 65+ years |
| HNO930 | M | 65+ years |
| HNO934 | F | 50-64 years |

**Supplemental Figure 1:**
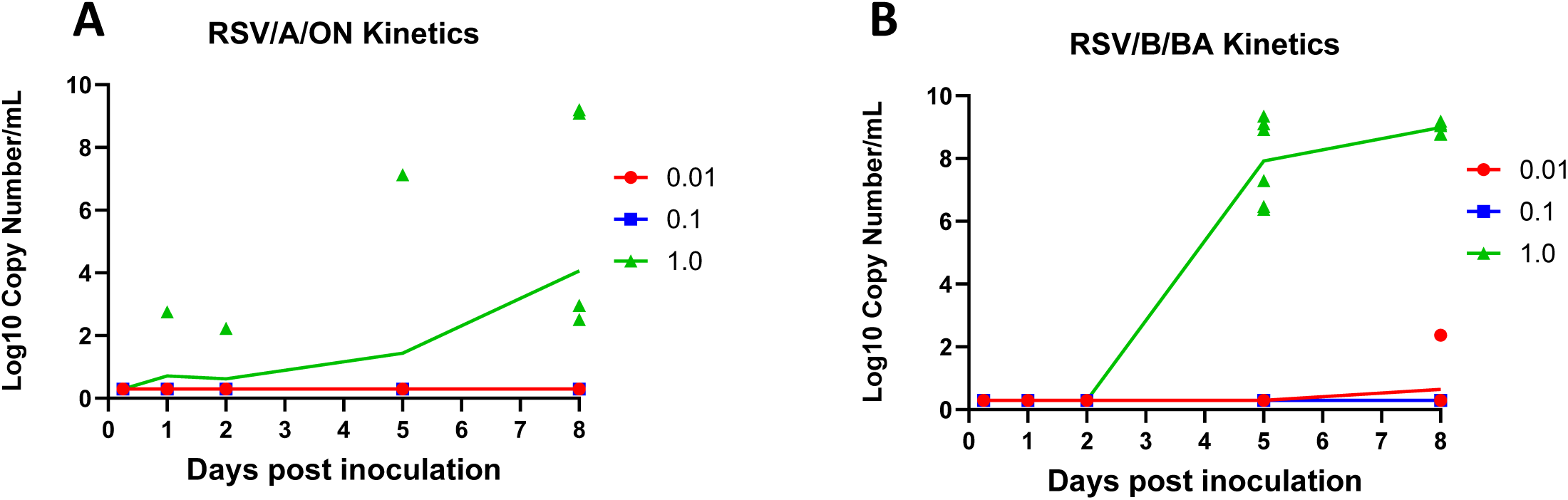
Viral RNA detection in apical lumen with increasing multiplicity of infection (MOI). qPCR of apical viral RNA with basolateral (A) RSV/A/ON and (B) RSV/B/BA inoculation at different MOIs. Data is pooled from one adult and one infant line.

**Supplemental Figure 2:**
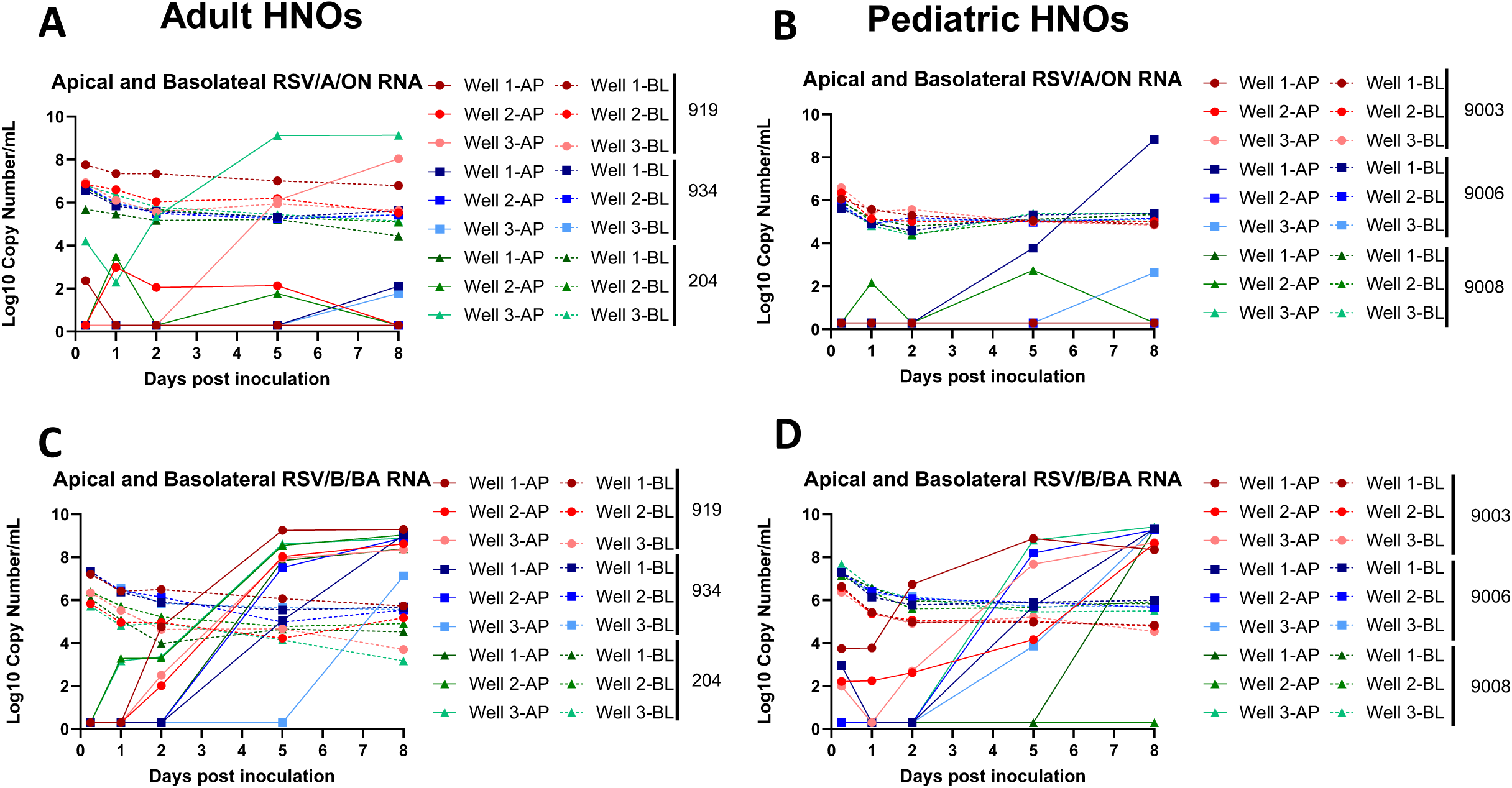
Viral RNA detection in basolateral media and apical lumen with basolateral inoculation of RSV/A/ON and RSV/B/BA. qPCR of apical and basolateral viral RNA with basolateral (A-B) RSV/A/ON and (C-D) RSV/B/BA inoculation in three technical replicates from three representative adult (919, 934, 204) lines (left) and three representative infant (right) HNO-ALI cultures (9003, 9006, 9008).

**Supplemental Figure 3:**
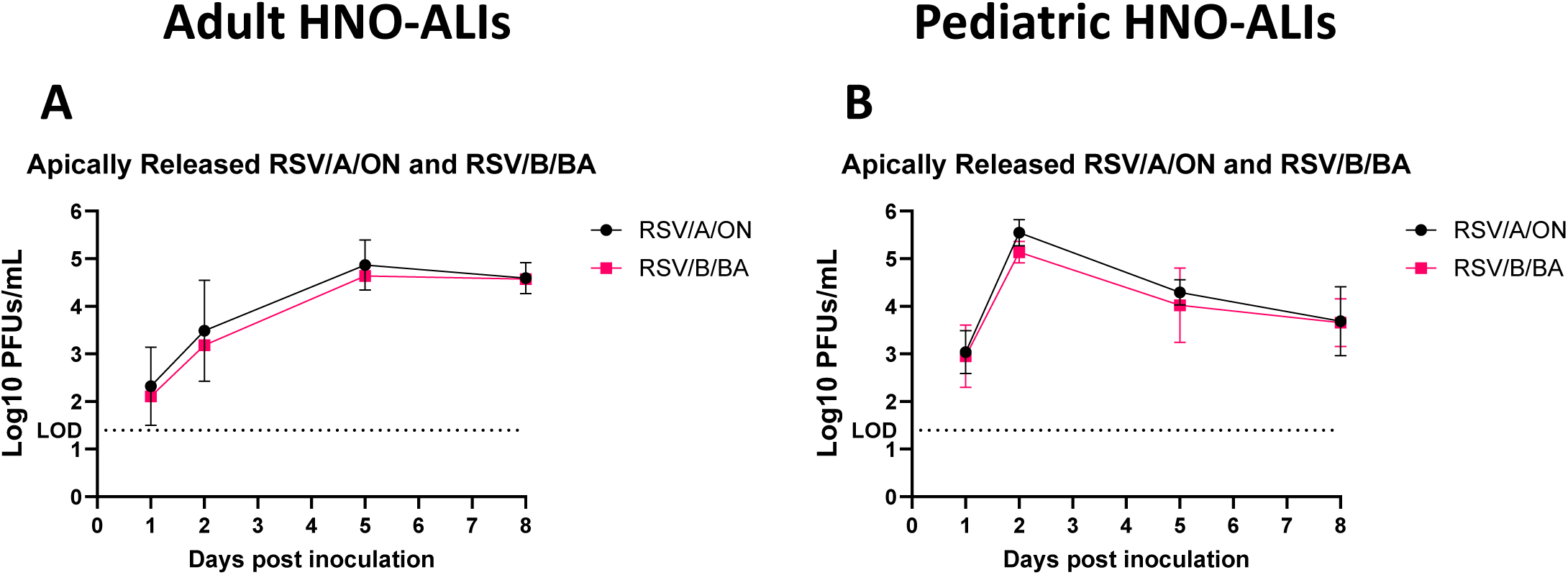
Viral kinetics of apical RSV/A/ON and RSV/B/BA infection. Pooled plaque assay data for (A) RSV/A/ON and (B) RSV/B/BA apical infection of adult (n=4 lines) and infant (n=4 lines) HNO-ALI cultures. LOD = limit of detection. Data represent mean ± standard deviation. Adapted from *Aloisio et al* (12).

**Supplemental Figure 4:**
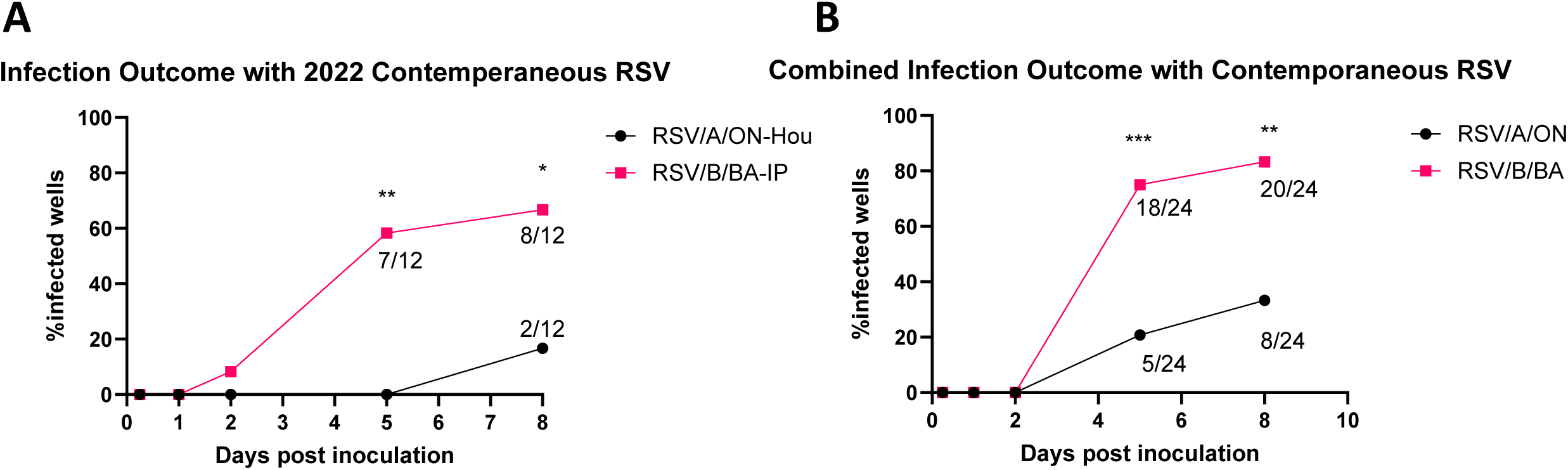
Infection outcome with additional contemporaneous RSV strains. (A) Total percentage of differentiated HNO-ALI transwells that demonstrated apical release of RSV with basolateral inoculation of contemporaneous strains of RSV/A/ON (RSV/A/ON-Hou) and RSV/B/BA (RSV/B/BA-IP) isolated in 2022 (n=4 adult lines, triplicate wells/line) (B) Combined infection outcome with two contemporaneous strains each of RSV/A/ON and RSV/B/BA strains (n=4 adult lines, triplicate wells/line). Statistical significance determined by Fisher’s Exact test; p values: *<0,05; **<0.01; *** <0.001

**Supplemental Figure 5:**
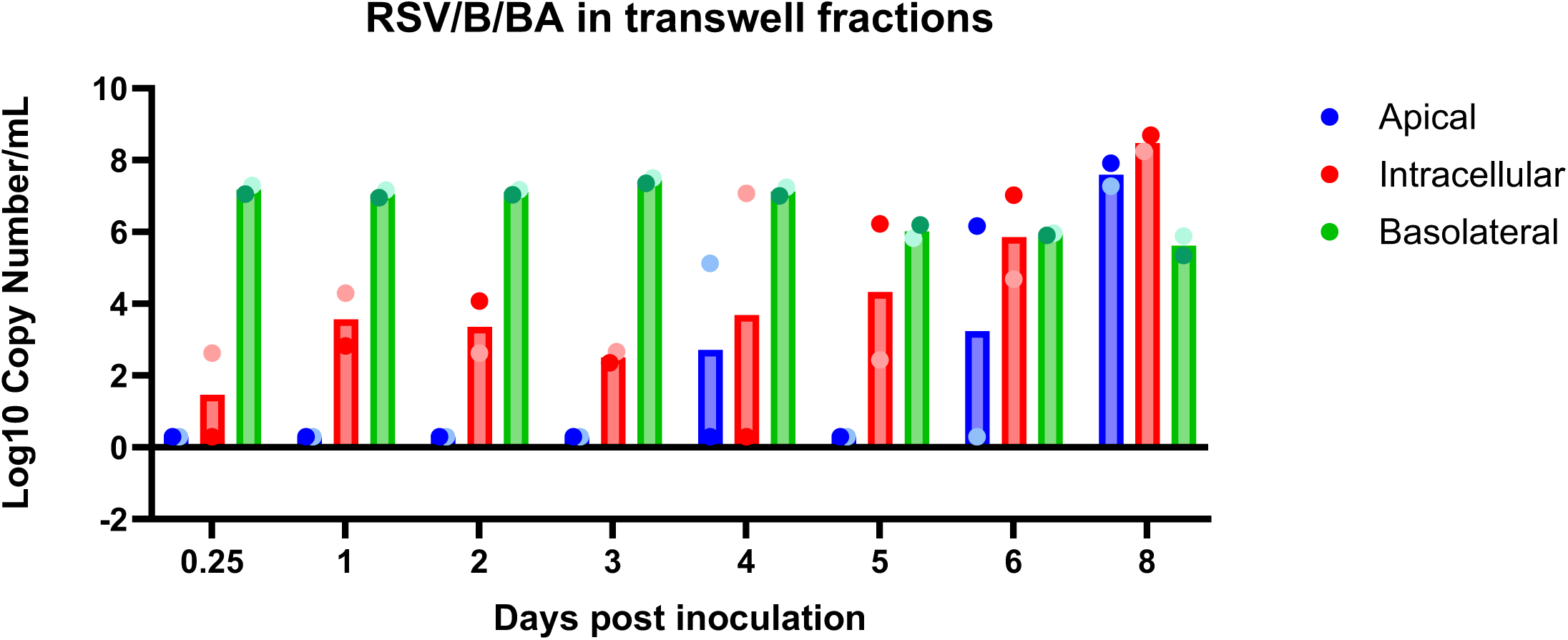
Viral RNA in apical, intracellular, and basolateral compartments. qPCR of apical, intracellular, and basolateral viral RNA with basolateral inoculation of RSV/B/BA in a single adult differentiated HNO-ALI line. Data represent mean of duplicate transwells/timepoint. Dark and light-colored data points are matched samples from the same transwell.

**Supplemental Figure 6:**
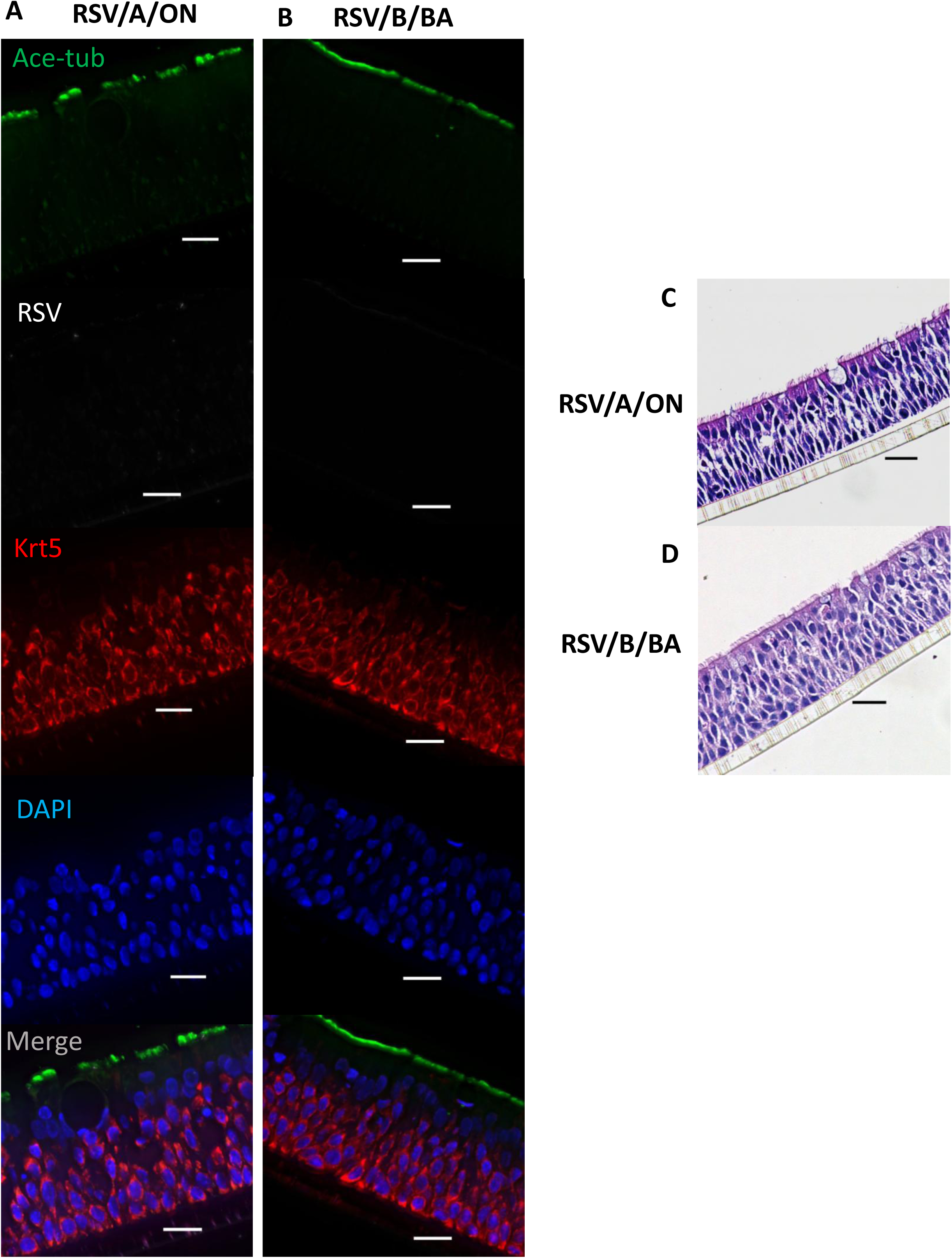
RSV/A/ON and RSV/B/BA basolateral inoculation of HNO-ALIs with no evidence of infection or epithelial damage. Representative IF images of a single representative infant HNO-ALI line basolaterally inoculated with (A) RSV/A/ON or (B) RSV/B/BA at 8 dpi. Ciliated cells – Acetylated alpha-tubulin ‘Ace-tub’ (green), RSV (white), Basal cells – Krt5(red), and nuclei – DAPI (blue). Representative H&E images of a single infant HNO-ALI line basolaterally inoculated with (C) RSV/A/ON or (D) RSV/B/BA at 8 dpi. Scale bar is 20 µm.

**Supplemental Figure 7:**
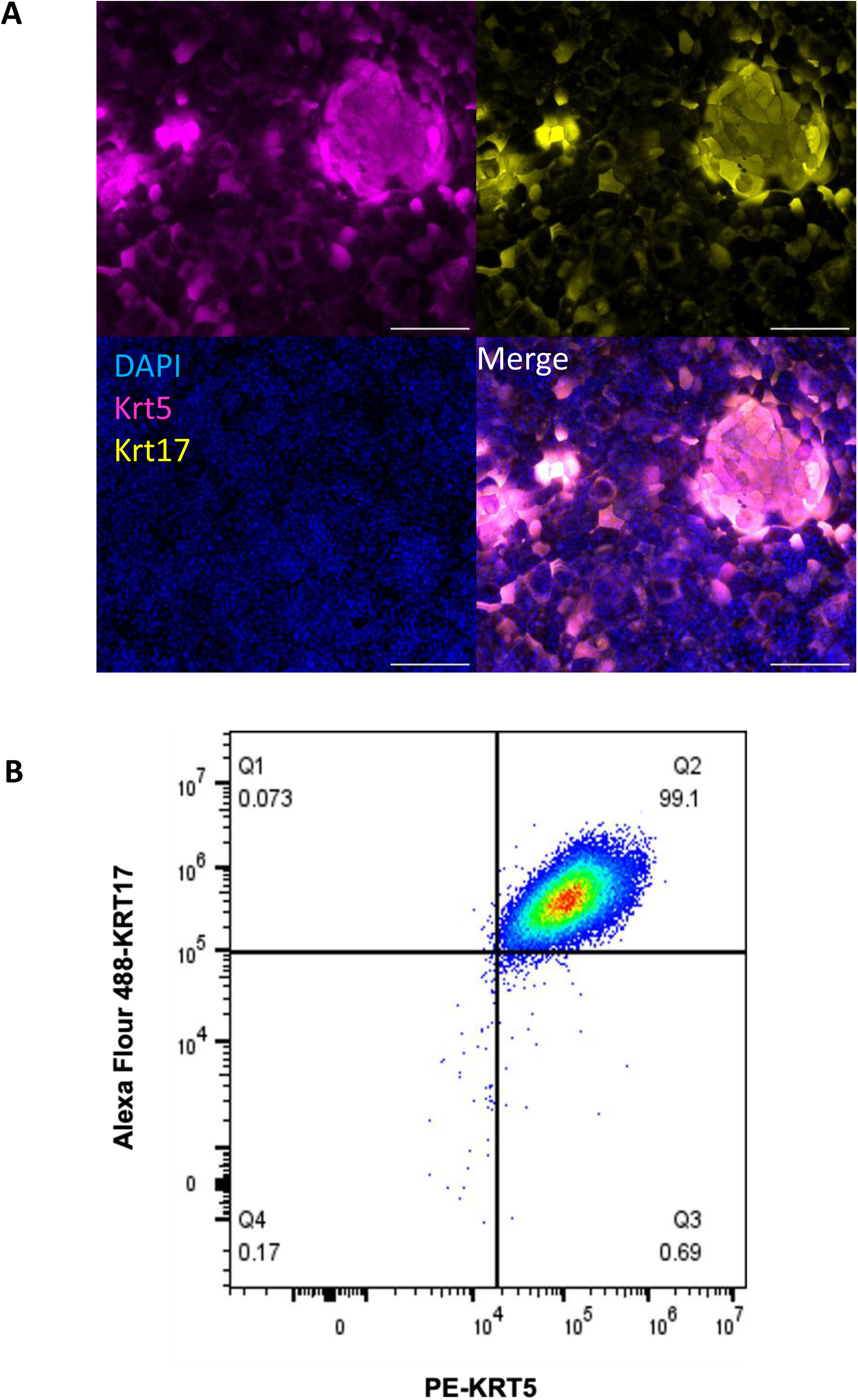
Pan-basal cell markers expressed in undifferentiated HNO-ALIs. (A) Representative image of an undifferentiated adult HNO-ALI stained for Krt5 (pink), Krt17 (yellow), and DAPI (blue). (B) Co-expression of Krt5 and Krt17 in a single undifferentiated adult HNO-ALI. Flow cytometric analysis of undifferentiated adult HNO-ALI cells stained for Krt5 (x-axis) and Krt17 (y-axis).

**Supplemental Figure 8:**
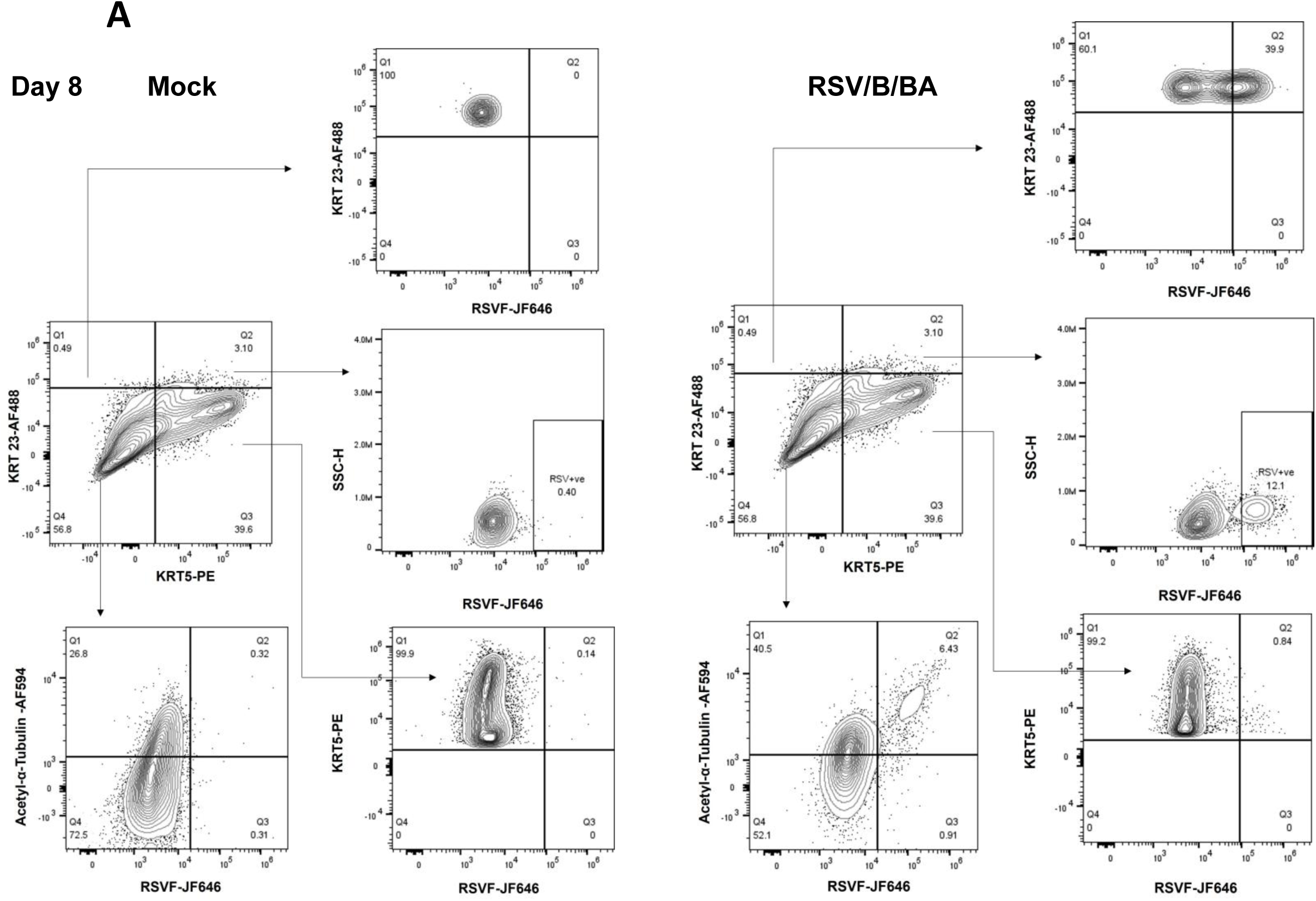
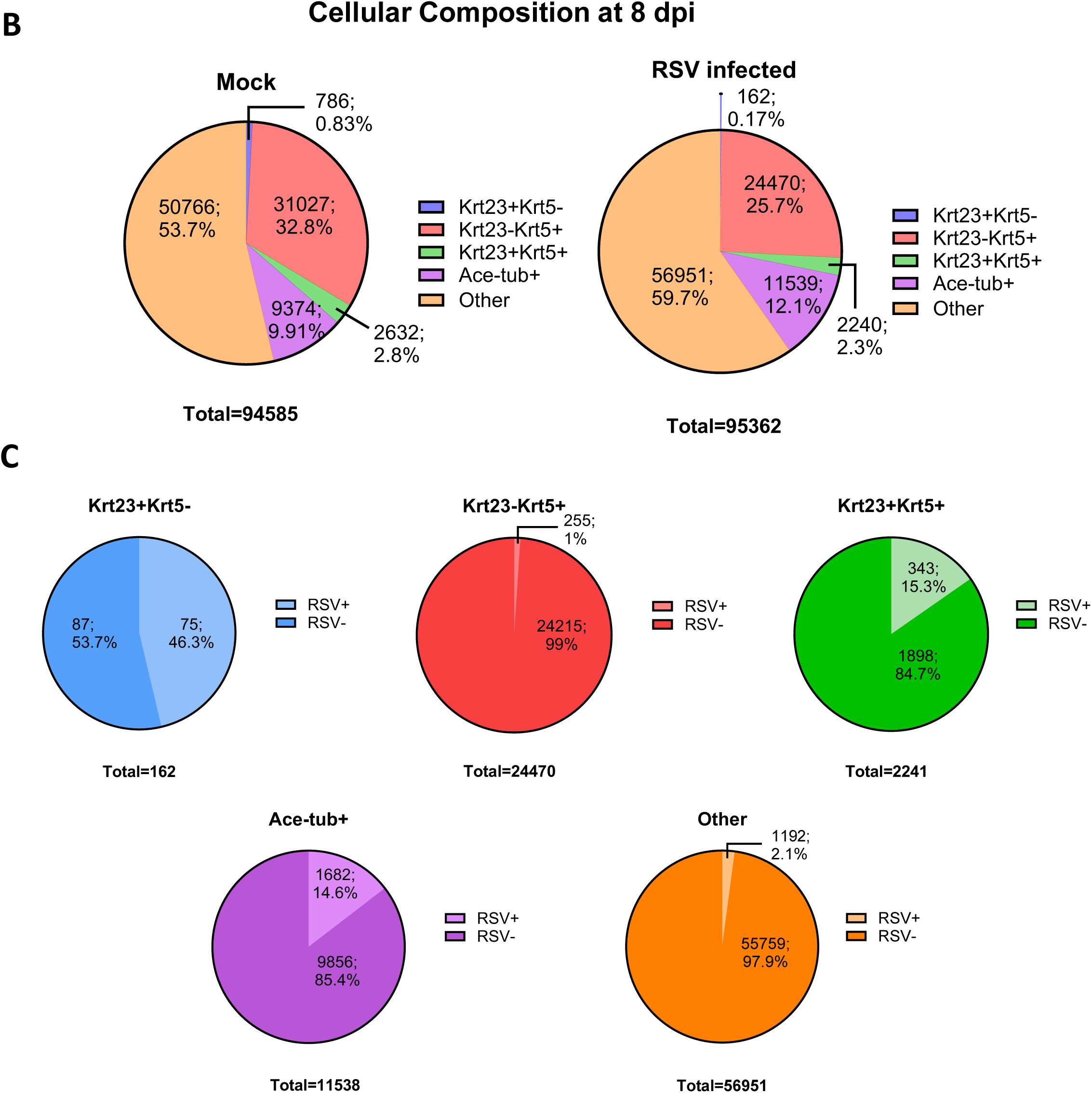
RSV selectively infects Krt23+ basal cells with basolateral inoculation of HNO-ALIs at 8 dpi. (A) Representative spectral flow cytometry plots showing gating strategy to identify Krt5+, Krt23+, and Ace-tub+ cell populations in basolateral mock and RSV/B/BA inoculated HNO-ALIs at 8 dpi. Further gating of RSV F protein was used to determine RSV infection in each cell population. (B) Pie charts summarizing the average proportion of each cell population in basolateral mock and RSV/B/BA inoculated HNO-ALIs at 8 dpi. (C) Pie charts summarizing RSV/B/BA infection in each cell population at 8 dpi.

